# Deep Learning Applications in Single-Cell Omics Data Analysis

**DOI:** 10.1101/2021.11.26.470166

**Authors:** Nafiseh Erfanian, A. Ali Heydari, Pablo Iañez, Afshin Derakhshani, Mohammad Ghasemigol, Mohsen Farahpour, Saeed Nasseri, Hossein Safarpour, Amirhossein Sahebkar

## Abstract

Traditional bulk sequencing methods are limited to measuring the average signal in a group of cells, potentially masking heterogeneity, and rare populations. The single-cell resolution, however, enhances our understanding of complex biological systems and diseases, such as cancer, the immune system, and chronic diseases. However, the single-cell technologies generate massive amounts of data that are often high-dimensional, sparse, and complex, thus making analysis with traditional computational approaches difficult and unfeasible. To tackle these challenges, many are turning to deep learning (DL) methods as potential alternatives to the conventional machine learning (ML) algorithms for single-cell studies. DL is a branch of ML capable of extracting high-level features from raw inputs in multiple stages. Compared to traditional ML, DL models have provided significant improvements across many domains and applications. In this work, we examine DL applications in genomics, transcriptomics, spatial transcriptomics, and multi-omics integration, and address whether DL techniques will prove to be advantageous or if the single-cell omics domain poses unique challenges. Through a systematic literature review, we find that DL has not yet revolutionized or addressed the most pressing challenges of the single-cell omics field. However, using DL models for single-cell omics has shown promising results (in many cases outperforming the previous state-of-the-art models) in data preprocessing and downstream analysis, but many DL models still lack the needed biological interpretability. Although developments of DL algorithms for single-cell omics have generally been gradual, recent advances reveal that DL can offer valuable resources in fast-tracking and advancing research in single-cell.

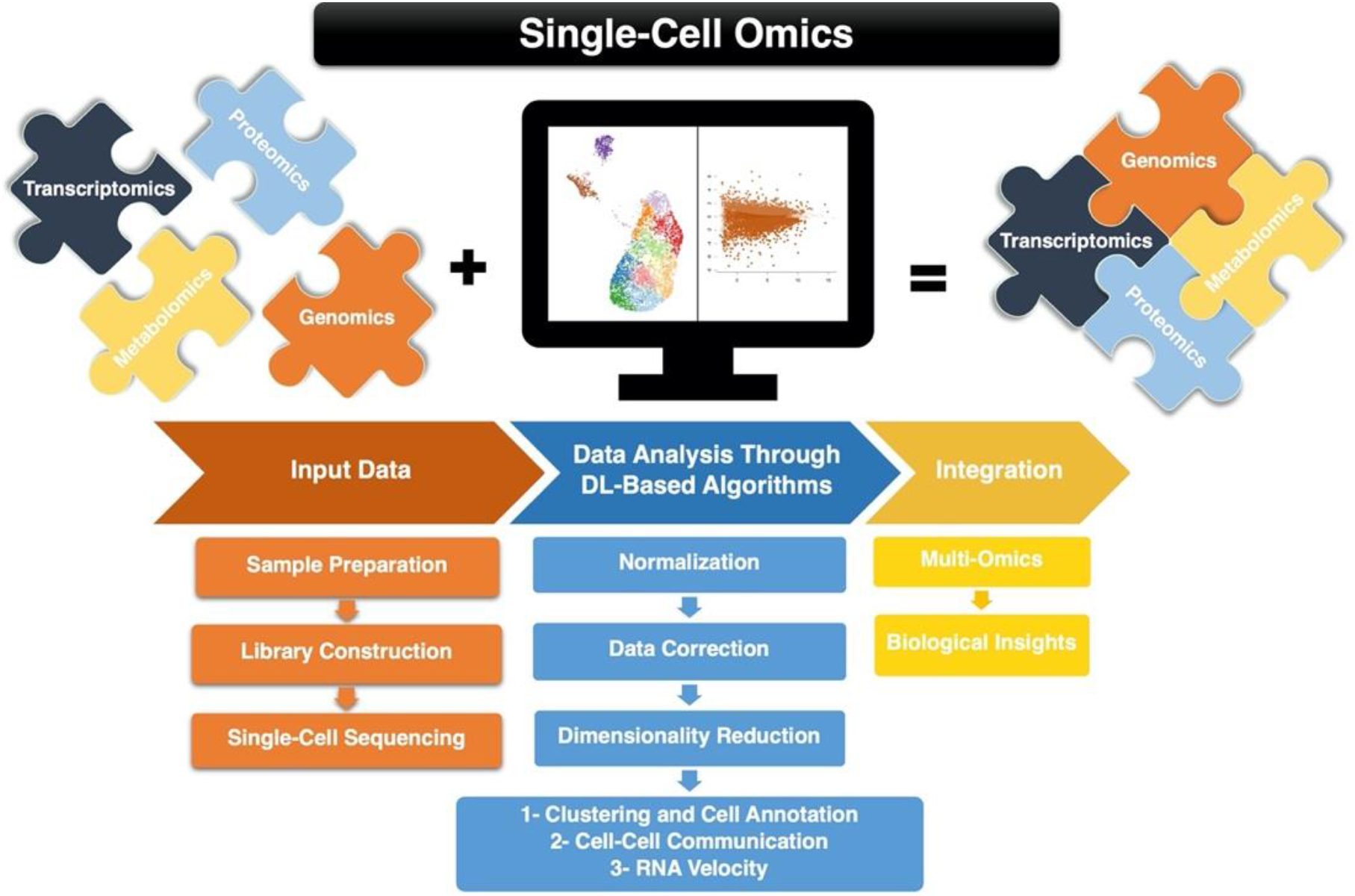

## 1. Introduction

Since single-cell sequencing (sc-seq) was highlighted as the “Method of the Year” in 2013 (Fritzsch, Dusny et al. 2012), sequencing individual cells at the single-cell resolution has become the norm for studying cell-to-cell heterogeneity. RNA and DNA single-cell measurements, and recently, epigenetic and protein levels, stratify cells at the highest possible resolution. That is, single-cell RNA sequencing (scRNA-seq) makes it possible to measure transcriptome-wide gene expression at the single-cell level. Such resolution enables researchers to distinguish different cell types based on their characteristics [see (Anchang, Hart et al. 2016, Haber, Biton et al. 2017, Tabula Muris, Overall et al. 2018, Han, Zhou et al. 2020)], organize cell populations, and identify cells transitioning between states. Such analyses provide a much better picture of tissues and the underlying dynamics in the development of an organism, which in turn allow for delineating intra-population heterogeneities that had previously been perceived as homogeneous by bulk RNA sequencing. Similarly, single-cell DNA sequencing (scDNA-seq) studies can reveal somatic clonal structures [e.g., in cancer, see (Roth, Khattra et al. 2014, Zafar, Navin et al. 2019)], thereby helping to monitor cell lineage development and providing insight into evolutionary mechanisms that function on somatic mutations.

The prospects resulting from sc-seq are tremendous: it is now possible to re-evaluate hypotheses on differences between predefined sample groups at a single-cell level regardless of samples being disease subtypes, treatment groups, or morphologically distinct cell types. As a result, in recent years, enthusiasm about the possibility of screening the basic units of life’s genetic material has continued to expand. Human Cell Atlas (Regev, Teichmann et al. 2017) is a prominent example: an effort to sequence the various cell types and cellular states that make up a human being. Encouraged by the great potential of single-cell investigation of DNA and RNA, there has been substantial growth in the development of related experimental technologies. In particular, the advent of microfluidics techniques and combinatorial indexing strategies (Hosokawa, Nishikawa et al. 2017, Zilionis, Nainys et al. 2017) has resulted in routinely sequencing hundreds of thousands of cells in a single experiment. This growth has also allowed a recent publication to analyze millions of cells at once (Cao, Spielmann et al. 2019). More large-scale sc-seq datasets are becoming accessible worldwide (with tens of thousands of cells), constituting a data explosion in single-cell analysis platforms. The continuous growth of the scale and quantity of available sc-seq data has raised significant questions: 1) How do we correctly interpret and analyze the increasing complexity of sc-seq datasets? 2) How can different types of data sets (mentioned above) provide a deeper understanding of the underlying biological dynamics for a specific condition? and 3) how can the acquired information be changed into practical applications in medicine, ranging from rapid and precise diagnoses to accurate medicine and targeted preventions. As we face the challenges of a rise in chronic diseases, aging populations, and limited resources, a transformation towards intelligent analysis, interpretation, and understanding of complex data is essential. In this paradigm shift, the rapidly emerging field of ML is central (Zheng and Wang 2019).

ML is the study of models which can learn from data without the need for an explicit set of instructions. Simultaneously with biomedical advancements of the past decade, there has been a surge in the development and application of ML algorithms, prominently led by advances in DL. Some of the earliest DL algorithms developed were intended to computationally model our brains’ learning process, therefore being called “Artificial Neural Networks” (ANN). DL models often consist of many processing layers (with many nodes in each layer), which enable them to learn a representation of data with several levels of abstraction. The recent improvements in computational hardware have made training DL models feasible, resulting in successful and revolutionary applications of such models across many domains.

This review aims to discuss DL applications in sc-seq analysis and elaborate on their instrumental role in improving sc-seq data processing and analysis. We first introduce some key concepts in DL, which enable us to discuss their applications in the sc-seq field. Because of the high technical noise and complexity of sc-seq data, we review appropriate techniques for analyzing such data, which ensure robustness and reproducibility of results. Finally, we conclude this work by discussing possible developments, highlighting both emerging obstacles and opportunities to advance the rapidly evolving field of single-cell omics.

## 2. Essential Concepts in Deep Learning

### 2.1 Feed Forward Neural Network

Due to the nature of sc-seq data, Feed Forward Neural Networks (FFNNs) are the most common architecture used as the core of many existing models for single-cell omics. FFNNs, the quintessential example of Artificial Neural Networks (ANNs), are composed of interconnected nodes (“artificial neurons”) that resemble and mimic the brain’s neuronal functions. The connections between the nodes (“edges”) strengthen or weaken as the learning progresses. Figure 1(a) illustrates an FFNN with an input layer (which senses and detects signals within the environment), three hidden layers (that process the signal sent by the input layer), and an output layer (the response to a signal or stimulus). FNNN can be viewed as a function that maps inputs from sc-seq space to an output space, often with fewer dimensions than the input space. This mapping is composed of simpler functions, each of which provides a different mathematical representation of the input. The true success of FFNN was realized when the capacity of such models was increased through the use of non-linear activation layers (such as Rectified Linear Units (Nair and Hinton 2010) and Hyperbolic Tangent) and adding more layers (depth).

**Figure 1.**
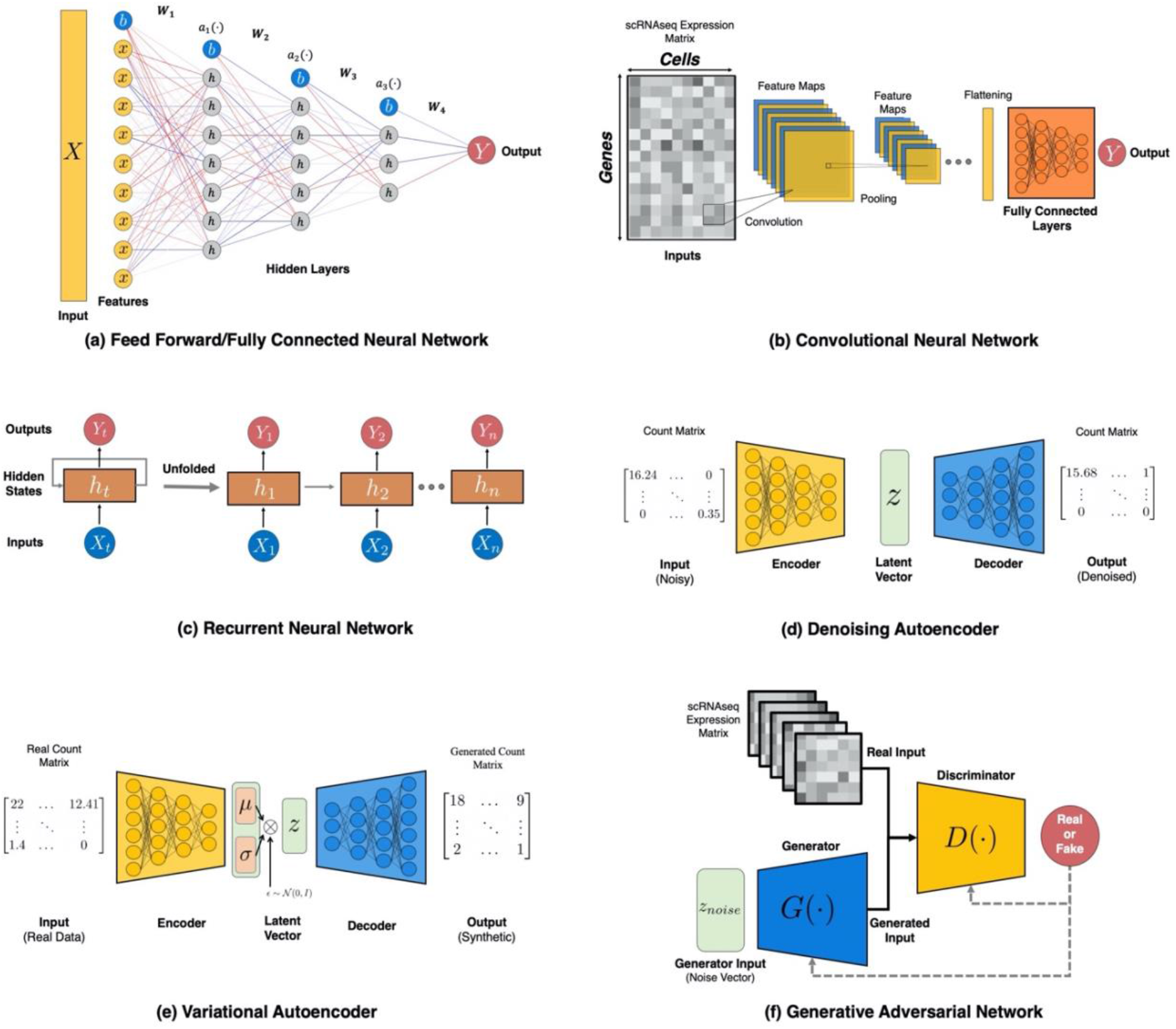
**(a)** An example of an FFNN architecture. In this network, the raw input is shown as *X*, the weight tensors are denoted *W*_*i*_, activation layers are α_*i*_ and the bias nodes are shown in blue nodes marked by *b*. The positive and negative weights are shown as blue and red edges, respectively, where a smaller numerical value corresponds to more transparent edges and vice versa. **(b)** An example of CNN architecture for classification. In this hypothetical architecture, the model feeds the inputs through the three stages of a CNN (with non-linear activation not depicted) to extract features. As is often the case, the output of the CNN is inputted to a fully connected neural network for classification. **(c)** An RNN depicted with its training flow, and an unrolled version showing the timestep-dependent inputs, hidden state, and outputs are marked. **(d)** An illustration of a denoising autoencoder. Autoencoders (AE) have an encoder network, which tries to extract the most critical features and map them to a latent space (often of a much smaller dimension than the input data). The decoder network takes in the latent vector produced by the encoder (denoted as z) and maps this representation back to the original input dimension. **(e)** The architecture of a Variational AE (VAE) that aims at generating synthetic scRNA-seq data using the gene expression values. **(f)** A depiction of generative adversarial network for generating in-silico scRNA-seq data. In this illustration, the scRNA-seq count matrix is converted to images that are used as real samples.

These architectural choices allow neural networks to approximate highly non-linear and complex functions (theoretically being “universal approximators”).

ANNs learn in the same manner as humans: by interacting with, and responding to, various stimuli within a local environment. FFNNs consist of numerous internal adjustable parameters (often on the order of hundreds of millions) that determine the input-output function of the machine. The objective of FFNNs (and all ANNs) is to minimize a loss function that teaches the network to perform specific tasks. This optimization is done via learning an optimal set of parameters (weights) that minimize the objective function. In FFNNs, all neurons are fully connected; thus, the model must learn the optimal contribution of each node, which determine the final output of the model. That is, the direct connection between any two nodes can have positive or negative weights (refer to Fig. 1(a) for a visualization of this concept). The output values are sequentially computed within the network’s hidden layers, with each input vector taking in the previous layer’s output until reaching the final output layer (this process is known as the forward propagation). This multi-stage propagation of features through the layers results in an abstract data representation attentive to small details while ignoring irrelevant information (Lin, Yang et al. 2019).

Initially assigned random values, the model weights are optimized throughout the training based on the loss function. Conventionally, the model optimization is done through gradient descent performed backwards in the network, which consists of two components: a numerical algorithm for efficient computation of the chain rule for derivatives (backpropagation) and an optimizer (e.g., Stochastic Gradient Descent or Adam (Kingma and Ba 2015). The optimizer is an algorithm that performs gradient descent, while backpropagation is an algorithm for computing the expression for gradients during the backward pass of the model. Deep neural networks (DNNs) are a subset of ANNs with more hidden layers between the input and output layers (usually greater than or equal to three hidden layers). This depth enables the model to learn complex nonlinear functions and allows the machine to learn a multi-step computer program (Goodfellow, Bengio et al. 2016). Several studies have shown that an increase in the network’s capacity (e.g., through depth or non-linearity) has resulted in better performance in recognition and prediction tasks (Nair and Hinton 2010, Sun, Liang et al. 2015, Bahdanau, Chorowski et al. 2016).

### 2.2 Common Algorithms in Deep Learning

In the following sections, we provide an overview of several frequently-used DL models for single-cell applications.

#### 2.2.1 Recurrent Neural Network (RNNs)

RNNs (Rumelhart, Hinton et al. 1986) are used for processing sequential data, including natural language and time series. RNNs process sequential inputs one at a time and implicitly maintain a history of previous elements in the input sequence. A typical architecture for RNNs is presented in Fig 1(c). Similar to FFNNs, RNNs learn by propagating the gradients of each hidden state’s inputs at discrete times. This process becomes more intuitive if we consider the outputs of the hidden units at various time iterations as if they were the outputs of different neurons in a deep multilayer network.

Due to the sequential nature of RNNs, the backpropagation of gradients would shrink or grow at each time step, causing the gradients to vanish or blow up, which makes RNNs notoriously hard to train. However, when these issues are averted (via gradient clipping or other techniques), RNNs are powerful models and gain state-of-the-art capabilities in many domains, such as the process of natural language. The training challenges combined with the nature of sc-seq data have resulted in fewer developments of RNNs for single-cell analysis. However, recently some studies have used RNNs and Long Short-Term Memory (a variant of RNNs) used for predicting cell types and cell motility (Hubel and Wiesel 1962, Kimmel, Brack et al. 2019).

#### 2.2.2 Convolutional Neural Network (CNNs)

CNNs (LeCun and Bengio 1995) are specialized types of networks that use convolution (the mathematical operation) instead of tensor multiplication (which is done in FFNNs) in at least one of their layers. This convolution operation makes CNNs ideal for processing data with a grid-like topology (images are the quintessential example of such datasets). Compared to other ANNs, CNNs have three key benefits: (i) sparse interactions, (ii) shared weights, and (iii) equivariant representations (LeCun and Bengio 1995). CNNs have been effectively used in many applications in computer vision and time-series analysis but are not utilized as frequently for sc-seq applications (since sc-seq datasets do not have a grid-like structure). However, some studies, such as Xu et al. (Xu, Zhang et al. 2020) have used CNNs after converting sc-seq data to images, which have shown promising results.

The architecture of CNNs was inspired by the organization of the visual cortex (LGN–V1–V2–V4–IT) hierarchy in the visual cortex ventral pathway (Hubel and Wiesel 1962), with the connectivity pattern trying to resemble the neural connections of our brains. A typical CNN architectural block is composed of a sequence of layers (usually three) which include a convolutional layer (affine transform), a detector stage (non-linear transformation), and a pooling layer. The learning unit of a convolutional layer is called a filter or kernel. Each convolutional filter is a matrix, typically of small dimensions (e.g. 3×3), composed of a set of weights that acts as an object detector and is continuously calibrated during the learning process. The goal of CNNs is to learn an optimal set of filters (weights) which can detected the needed features for a specific task (e.g. image classification). The result of the convolution between the input data and the filter’s weights is named a feature map. Once a feature map is available, each value of this map is passed through a non-linearity (e.g. ReLU). The output of a convolutional layer is made of as many stacked feature maps as the number of filters present within the layer. There are two key ideas behind this design: first, in data with grid-like topology (such as images), local neighbors have highly correlated information. Second, equivariance to translation can be obtained if units at different locations share weights. In other words, parameter sharing in CNNs allows for the detection of features regardless of the locations that they appear in. An example of this would be detecting a car. In a dataset, a car could appear at any position in a 2D image, but the network should be able to detect it regardless of the specific coordinates (Zhao, Zheng et al. 2019).

One way of achieving equivariance to translation is to utilize pooling layers. In the pooling layers, we use the outputs of the detector stage (at certain locations) to calculate a summary statistic for a rectangular window of values (e.g., calculating the mean of a 3×3 patch). There are many pooling operations, with the common ones being max-pooling (taking the maximum value of a rectangular neighborhood), mean-pooling (taking the average), and *L*_*2*_ norm (taking the norm). In all cases, rectangular patches from one or several feature maps are inputted to the pooling layer, where semantically similar features are merged into one. Pooling decreases the dimension of learned representations, and makes the model insensitive to small shifts and distortions (LeCun, Bengio et al. 2015). CNNs typically have an ensemble of stacked convolution layers, non-linearity, and pooling layers, followed by fully connected layers that produce the final output of the network. Fig. 1(b) illustrates an example of a typical CNN architecture. The backpropagation of gradients through CNNs is analogous to DNNs, enabling the model to learn an optimal set of filters.

#### 2.2.3 Autoencoders (AEs)

AEs are neural networks that aim to reconstruct (or copy) the original input via a non-trivial mapping. Conventional AEs have an “hour-glass” architecture with two mirroring networks: an encoder and a decoder. The encoder’s task is to map the input data to a latent space, often of a much smaller dimension than the original input space. The encoder is responsible for data compression and feature extraction, forming the narrowing part of the hourglass architecture (see Fig. 1(d)). The output of the encoder network (latent vector) will contain the most important features present in the data in a compressed form. Conversely, the decoder is tasked with mapping the latent vector back to the original input dimension and reconstructing the original data. In the ideal case, the decoder’s output will be an exact copy of the training sample.

AEs were traditionally used for dimensionality reduction and denoising, trained by minimizing a mean squared error (MSE) objective between the input data and the reconstructed sample (output of the decoder). Fig. 1(d) depicts an example of a denoising AE. Over time, the AE framework has been generalized to stochastic mappings of an encoder distribution and a decoder distribution. A well-known example of such generalization is Variational Autoencoders (VAEs) (Kingma and Ba 2015), where using the same hour-glass architecture, one can generate new samples drawn from an approximated posterior for an assumed prior distribution. Both traditional AEs and VAEs have practical applications in many biological fields.

#### 2.2.4 Variational Autoencoders (VAEs)

VAEs (Kingma and Ba 2015, Kingma and Welling 2019) are generative models that learn latent-variable and inference models simultaneously, i.e., they are made up of generative and inference models. VAEs are AEs that employ variational inference to recreate the original data, allowing them to produce new (or random) data that is “similar” to that which already exists in a dataset (illustrated in Fig. 1(e)). VAEs have better mathematical properties and training stability than Generative Adversarial Networks (GANs), but they suffer from two major weaknesses: classic VAEs create “blurry” samples (those that adhere to an average of the data points), rather than the sharp samples that GANs generate due to adversarial training. Introspective VAEs (IntroVAEs) Huang et al. have generally solved this issue by specifying an adversarial training between the encoder and the decoder (Huang, Li et al. 2018). IntroVAEs are single-stream generative algorithms that assess the quality of the images they generate. They have largely been employed in computer vision, where they have outperformed their GAN counterparts in applications like synthetic image generation (Huang, Li et al. 2018) and single-image super-resolution (Heydari and Mehmood 2020).

The other major issue with VAEs is *posterior collapse*: when the variational posterior and actual posterior are nearly identical to the prior (or collapse to the prior), which results in poor data generation quality (He, Spokoyny et al. 2019). Posterior collapse has been attributed to VAE’s distribution regularization term in the objective function (Lucas, Tucker et al. 2019), i.e. when the prior and posterior divergence is close to zero. Studies aimed at reducing posterior collapse can be divided into two categories: (i) solutions aimed at weakening the generative model (Semeniuta, Severyn et al. 2017, Yang, Hu et al. 2017), (ii) alterations to the training purpose (Tolstikhin, Bousquet et al. 2017, Yang, Hu et al. 2017, Zhao, Song et al. 2017), and (iii) alterations to the training procedure (He, Spokoyny et al. 2019, Heydari, Thompson et al. 2019). If both issues mentioned above can be addressed, VAEs have shown to perform comparable (or similar) to GANs while training faster due to the simpler training procedure.

#### 2.2.5 Generative Adversarial Networks (GANs)

GANs (Goodfellow, Pouget-Abadie et al. 2014) can generate realistic synthetic data and have been effectively utilized in a variety of computer vision tasks (Dziugaite, Roy et al. 2015, Vondrick, Pirsiavash et al. 2016, Zhu, Krähenbühl et al. 2016), natural language processing (Yang, Chen et al. 2017, Fedus, Goodfellow et al. 2018), time series synthesis (Esteban, Hyland et al. 2017, Engel, Agrawal et al. 2019), and bioinformatics (Marouf, Machart et al. 2020). GANs are made up of a generator network (G) and a discriminator network (D) that compete in a zero-sum game; we present the architecture of GANs in Fig. 1(f). The goal of the G network is to generate fake samples that resemble the distribution of the real data, “fooling” the D network into believing that these fake samples are real. Conversely, D trains to learn the difference between real and synthetic samples and “discriminate” between them. In each GAN training iteration, the entire system is re-adjusted to update both G and D parameters. In this process and through many iterations, the generator learns to make more realistic samples which deceive the discriminator as real data. At the same time, the discriminator is learning the distinction between real and generated data (from the G network). GANs’ ability to produce realistic samples is attributed to the adversarial training between G and D networks. Compared to other generative models, GANs have several advantages, such as the flexibility to learn any distribution, requiring no assumptions on the prior distribution, and no limitations on the size of the latent space.

Despite these advantages, GANs are notoriously difficult to train since achieving Nash equilibrium for G and D is extremely difficult (Wang, She et al. 2021). Another drawback of GANs is vanishing gradients, which occurs if D learns the distinction between real and generated data well too quickly, prohibiting G from training propery (Arjovsky, Chintala et al. 2017). Another problem with GANs is “mode collapse,” which occurs when G produces only a small number of outputs that potentially trick D. This happens when G has trained to map many noise vectors to the same output that D recognizes as real data. Quantifying how much GANs have learned the distribution of real data is often difficult, hence one of the most common methods of assessing GANs is to evaluate the output directly (Larsen, Sønderby et al. 2016), which could be laborious. Even though certain GAN variations have been proposed to reduce vanishing gradients and mode collapse, (e.g., Wasserstein-GANs (WGANs) (Arjovsky, Chintala et al. 2017) and Unrolled-GANs (Metz, Poole et al. 2016), the convergence of GANs is still a big issue.

## 3. DL Applications in SC Omics

### 3.1 Deep Learning in Single-Cell Transcriptomics

Single-cell RNA sequencing (scRNA-seq) has improved our understanding of biological processes substantially in recent years. Researchers have shown the potentials of single-cell (SC) transcriptomics through studying the cellular heterogeneity of many organisms, such as humans, mice, zebrafish, frogs, and planaria (Tabula Muris, Overall et al. 2018, Han, Zhou et al. 2020), and uncovering previously unknown cell populations (Montoro, Haber et al. 2018, Plass, Solana et al. 2018). Moreover, these studies have also highlighted the need for better computational methods which can facilitate the analysis of large and complex scRNA-seq datasets. In the following sections, we provide an overview of existing computational approaches for the various stages of scRNA-seq analysis (summarized in Fig. 2).

**Figure 2.**
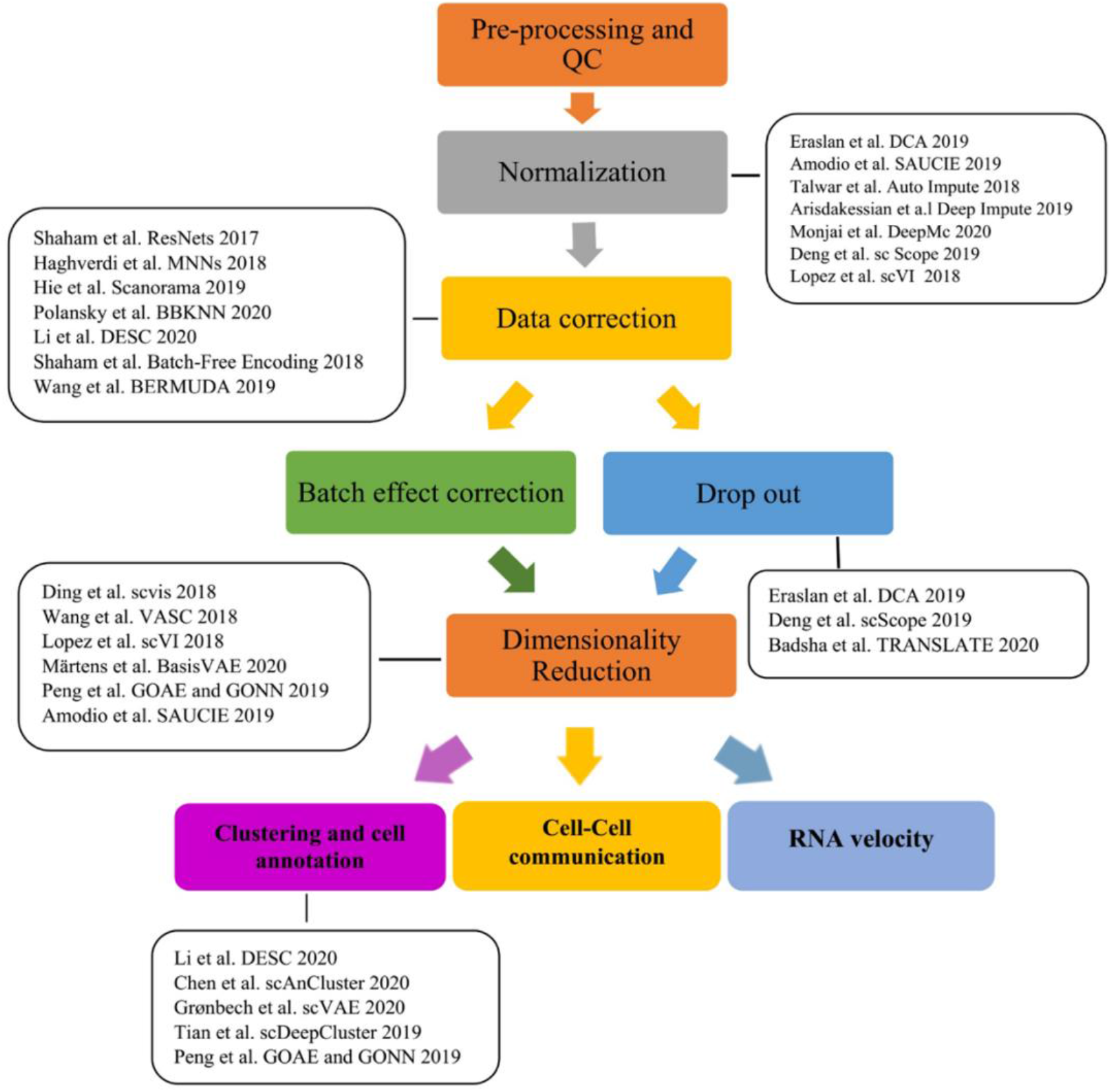
Workflow of RNA-seq data analysis

#### 3.1.1 Pre-processing and quality control

Cell barcodes are designed to delineate various cell populations present within a sequenced sample. However, barcodes can mistakenly tag several cells (doublet) or not tag any cells at all (empty droplet/well), which prompts the quality control (QC) step in scRNA-seq analysis. Many raw data pre-processing pipelines, including Cell Ranger (Zheng, Terry et al. 2017), indrops (Klein, Mazutis et al. 2015), SEQC (Azizi, Carr et al. 2018), and z-unique molecular identifiers (zUMIs) (Parekh, Ziegenhain et al. 2018), can perform QC. The dimension of the count matrix generated by sequencing technologies depends on the the number of barcodes and transcripts. Though the noise rate in measurements varies across reads and count data, standard research pipelines very often emply the same processing techniques (Lafzi, Moutinho et al. 2018).

Despite scRNA-seq data’s richness which can offer significant and deeper insights, the data’s complexity and noise are far higher than traditional bulk RNA-seq, making it challenging to process the raw data for downstream analysis. Unwanted variations such as biases, artifacts, etc., require extensive QC and normalization efforts (Jiang, Thomson et al. 2016). The number of counts per barcode (count depth), the number of genes per barcode, and the fraction of counts from mitochondrial genes per barcode are three QC covariates widely used in the QC step (Griffiths, Scialdone et al. 2018). On the other hand, other experimental factors (such as damaging the sample during dissociation) could result in low-quality scRNA-seq libraries, which can yield erroneous findings in downstream analyses. Currently, there is an unmet need for developing more efficient and accurate methods for filtering low-quality cells during the library preparation.

Given the limited number of studies regarding pre-processing and quality control, we will focus on the DL applications that are most related to normalization, data correction, and downstream analysis.

#### 3.1.2 Normalization

Normalization is a crucial first step in pre-processing scRNA-seq expression data to address the constraints caused by low input content, or the different types of systematic measuring biases (Bacher, Chu et al. 2017). Normalization aims to detect and remove changes in measurements between samples and features (e.g., genes) caused by technical artifacts or unintended biological effects (e.g., batch effects) (Hogan, Courtier et al. 2019). Methods designed for normalizing bulk RNA-seq and microarray data are often used to normalize scRNA-seq data. However, these techniques often ignore essential aspects of scRNA-seq results (Hogan, Courtier et al. 2019). For scRNA-seq data, a few families of normalization techniques have been developed, such as scaling techniques (Lun, Bach et al. 2016), regression-based techniques for identified nuisance factors (Buettner, Natarajan et al. 2015, Bacher, Chu et al. 2017), and techniques based on spike-in sequences from the External RNA Controls Consortium (ERCC) (Ding, Zheng et al. 2015, Vallejos, Marioni et al. 2015). However, these methods are specific to certain experiments and can not be applied to all research designs and experimental protocols. Although some DL-based methods have been proposed to generalize the normalization stage, accounting for technical noise of scRNA-seq data still remains a challenge and an active area of research within the field (Zheng and Wang 2019).

#### 3.1.3 Data correction

Although normalization aims to address the noise and bias in the data, normalized data can still contain unexpected variability. These additional technical and biological variables, such as batch, dropout and cell cycle effects are accounted for during the “data correction” stage, which depends on the downstream analysis (Luecken and Theis 2019). In addition, it is a recommended practice to address biological and technical covariates separately (Luecken and Theis 2019), since they serve different purposes. Given the mentioned subtleties, designing DL models that can address most of these challenges is difficult. Therefore, there are currently no DL models that are widely used for data correction within the field.

##### 3.1.3.1 Dropout

Compared to bulk RNA-seq, scRNA-seq datasets are noisy and sparse and pose unique nuances such as “dropout” [which is one of the most significant issues in this field (Kharchenko, Silberstein et al. 2014, Gong, Kwak et al. 2018)]. Dropout occurs when a gene is observed at a moderate or high expression level in one cell but is not detected in another cell (Qiu 2020). Dropout events can occur during library preparation (e.g. extremely low levels of mRNA in single cells) or due to biological properties (the stochastic aspect of gene expression in multiple cells)(Ran, Zhang et al. 2019). Additionally, shorter genes have lower counts and a greater dropout rate (Zappia, Phipson et al. 2017). Overall, a low RNA capture rate results in the inability of detecting an expressed gene, leading to a “false” zero, known as a dropout event. Furthermore, it has been suggested that sometimes near-zero expression measurements can also be dropouts (Lin, Troup et al. 2017). Dropout events will introduce technical variability and noise, adding an extra layer of difficulty in analyzing scRNA-seq (Sengupta, Rayan et al. 2016), and downstream analyses such as clustering and pseudo-time reconstruction (Arisdakessian, Poirion et al. 2019).

It is essential to understand the difference between “false” and “true” zero counts. True zero counts mean that a gene is not expressed in a particular cell type, indicating true cell-type-specific expression (Eraslan, Simon et al. 2019). Hence, it is important to note that zeros in scRNA-seq data do not necessarily translate to missing values and must remain in the data. However, the false zeros (missing values) must be imputed to further improve the analysis. The missing values are replaced with either random values or by an imputation method (Eraslan, Simon et al. 2019). Imputation approaches designed for bulk RNA-seq data may not be suitable for scRNA-seq data for multiple reasons, mainly due to scRNA-seq’s heterogeneity and dropouts. ScRNA-seq has much higher cell-level heterogeneity than bulk RNA-seq data; scRNA-seq has cell-level gene expression data while bulk RNA-seq data represents the averaged gene expression of the cell population. Additionally, the number of missing values in bulk RNA-seq data is much lower compared to scRNA-seq (Gong, Kwak et al. 2018). Given these factors and the non-trivial difference between true and false zero counts, classic imputation approaches with specified missing values are often not appropriate for scRNA-seq data, and scRNA-seq-specific dropout imputation methods are required.

Current scRNA-seq imputation methods can be divided into two groups: (i) those that change all gene expression levels, such as Markov Affinity-based Graph Imputation of Cells (MAGIC) (van Dijk, Nainys et al. 2017) and single-cell analysis via expression recovery (SAVER) (Huang, Wang et al. 2018), and (ii) methods that impute drop-out events (zero or near-zero counts) alone, such as scImpute (Li and Li 2018), DrImpute (Gong, Kwak et al. 2018), and LSImpute (Moussa and Măndoiu 2019). These techniques can fail to account for the non-linearity of the data’s count structure. Moreover, as larger scRNA-seq datasets become available and common, imputation methods should scale to millions of cells. However, many of the earlier models are either incapable of or very slow at processing datasets of larger size (tens of thousands or more) (Eraslan, Simon et al. 2019). As a result, many have resorted to designing DL-based approaches to combat these challenges, both on the technical and efficiency fronts.

Most of DL algorithms for imputing drop-out events are based on AEs. For example, in 2018, Talwar et al. proposed AutoImpute, a technique for retrieving the whole gene expression matrix using overcomplete AEs to impute the dropouts. AutoImpute learns the underlying distribution of the input scRNA-seq data and imputes missing values based on the learned distribution, with minor modifications to the biologically silent gene expression values. Through expanding the expression profiles into a high-dimensional latent space, AutoImpute learns the underlying distribution and patterns of gene expression in single cells and reconstructs an imputed model of the expression matrix. At the time, Talwar et al. claimed that their system was the only model that could perform imputation on the largest of the nine datasets they studied (68K PBMC, which contains ∼68,000 cells), without running out of memory (Talwar, Mongia et al. 2018).

In another study, Eraslan et al. proposed the Deep Count AE network (DCA). DCA uses a negative binomial noise model both with and without zero-inflation to account for the count distribution and overdispersion while capturing nonlinear gene-gene dependencies. Since their approach scales linearly with the number of cells, DCA can be used on datasets with millions of cells. DCA also depends on gene similarities; using simulated and true datasets, DCA denoising enhances several traditional scRNA-seq data analyses. One of the key benefits of DCA is that it only requires the user to define the noise model. Current scRNA-seq approaches depend on a variety of hypotheses and often use standard count distributions, such as zero-inflated negative binomial. DCA increases biological exploration by outperforming current data imputation approaches in terms of quality and time. Overall, DCA calculates the “dropout probability” of a zero-expression value due to scRNA-seq dropout and imputes the zeros only when the probability is high. Consequently, while DCA effectively detects true zeros, it can be biased when dealing with nonzero values (Eraslan, Simon et al. 2019).

Badsha et al. propose TRANSfer learning with LATE (TRANSLATE) (Badsha, Li et al. 2020), a DL model for computing zeros in scRNA-seq datasets which are extremely sparse. Their nonparametric approach is based on AEs and builds on their previous method, Learning with AuToEncoder (LATE). The key presumption in LATE and TRANSLATE is that all zeros in the scRNA-seq data are missing values. In most cases, their approach achieves lower mean squared error, restores nonlinear gene-gene interactions, and allows for improved cell type separation. Both LATE and TRANSLATE are also very scalable, and when using a GPU, they can train on over a million cells in a few hours. TRANSLATE has shown better performance on inferring technical zeros than other techniques, while DCA is better at inferring biological zeros than TRANSLATE.

Sparse Autoencoder for Unsupervised Clustering, Imputation, and Embedding (SAUCIE) (Amodio, Van Dijk et al. 2019) is a regularized AE that denoises and imputes data using the reconstructed signal from the AE. Despite the noise in the input data, SAUCIE can restore the significant relationships across genes, leading to better expression profiles which can improve downstream analyses such as differential gene expression (Amodio, Van Dijk et al. 2019).

ScScope (Deng, Bao et al. 2019) is a recurrent AE network that iteratively handles imputation by employing a recurrent network layer; taking the time recurrence of ScScope to one (i.e. *T=1)* will reduce the model to a traditional AE. Given that ScScope is a modification of traditional AEs, its runtime is similar to other AE-based models (Deng, Bao et al. 2019).

A few non-AE-based models have also been developed for imputation and denoising of scRNA-seq data. DeepImpute (Arisdakessian, Poirion et al. 2019) uses several sub-neural networks to impute groups of target genes using signals (genes) that are strongly associated with the target genes. Arisdakessian et al. demonstrate that DeepImpute has a better performance than DCA, contributing the advantages to their divide-and-conquer approach (Arisdakessian, Poirion et al. 2019).

Mongia et al. (Mongia, Sengupta et al. 2020) introduced Deep Matrix Completion (deepMC), an imputation method based on deep matrix factorization for missing values in scRNA-seq data that utilizes a feed backward neural network. In most of their experiments, deepMC outperformed other existing imputation methods while not requiring any assumption on the prior distribution for the gene expression. We predict that deepMC will be the preferred initial approach for imputing scRNA-seq data, given the superior performance and simplicity of the model.

Single-cell variational inference (scVI) is another DNN algorithm introduced by Lopez et al (Lopez, Regier et al. 2018). ScVI is based on a hierarchical Bayesian model and uses a DNN to define the conditional probabilities, assuming either a negative binomial or a zero-inflated negative binomial distribution (Lopez, Regier et al. 2018). Lopez et al. show that scVI can accurately recover gene expression signals and impute the zero-valued entries, potentially enhancing the downstream analyses without adding any artifacts or false signals.

Recently, Patruno et al. (Patruno, Maspero et al. 2020) compared 19 denoising and imputation methods, based on numerous experimental scenarios such as recovery of true expression profiles, characterization of cell similarity, identification of differentially expressed genes, and computation time. Their results showed that ENHANCE (Expression deNoising Heuristic using Aggregation of Neighbors and principal Component Extraction), MAGIC, SAVER, and SAVER-X offer the best overall results when considering efficiency, accuracy and robustness for the investigated tasks (Patruno, Maspero et al. 2020).

It is important to note that traditional methods, despite their current success, are not well-suited for large-scale scRNA-seq studies. As larger scRNA-seq datasets become the norm, we anticipate that DL-based models will prove to be advantageous. Therefore, more work is required to build upon the existing DL methods for imputing dropout effects and better managing technical zeros while retaining biological zeros.

##### 3.1.3.2 Batch effects correction

When samples are conducted in separate batches, the term ‘batch effect” is used to describe the variation caused by technical effects. Different types of sequencing machines or experimental platforms, laboratory environments, different sample sources, and even technicians who perform the experiments can cause batch effects (Fei and Yu 2020). Removing and accounting for batch effects is often helpful and recommended, however, the success varies significantly across different studies. For example, batch effect removal on bulk RNA-seq data from Encyclopedia of DNA Elements (ENCODE) human and mouse tissues (Lin, Lin et al. 2014) is a recommended standard data preparation step. Batch effect correction has been an active area of research since the microarray time. Johnson et al. suggested parametric and non-parametric empirical Bayes frameworks for adjusting the data for batch effects removal (Johnson, Li et al. 2007). In recent years and with an increased level of complexity in sequencing datasets, more involved batch effect correction methods have been proposed and used (Fei and Yu 2020). However, a majority of the existing approaches require biological group expertise for each observation and were originally designed for bulk or microarray RNA-seq data. Given the heterogeneity present within scRNA-seq date, these earlier techniques are not well suited for single-cell analysis in certain cases (Luo and Wei 2019). Batch effects in scRNA-seq data may have a substantial impact on downstream data analysis, impacting the accuracy of biological measurements and ultimately contributing to erroneous conclusions (Büttner, Miao et al. 2019). Therefore, alternative batch effect correction techniques for scRNA-seq data have been developed to address the specific needs of single-cell datasets.

Several statistical methods, including linear regression models like ComBat (Johnson, Li et al. 2007), and nonlinear models like Seurat’s canonical correlation analysis (CCA) (Zhang, Wu et al. 2019) or scBatch (Fei and Yu 2020), have been designed to eliminate or minimize scRNA-seq batch effects while aiming to maintain biological heterogeneity of scRNA-seq data. Additionally, some differential testing frameworks such as Linear Models for Microarray (limma) (Ritchie, Phipson et al. 2015), Model-based Analysis of Single-cell Transcriptomics (MAST) (Finak, McDavid et al. 2015), and DESeq2 (Love, Huber et al. 2014) already integrate the batch effect as a covariate in model design.

Haghverdi et al. (Haghverdi, Lun et al. 2018) developed a novel and efficient batch correction method for single-cell data which detects cell mappings between datasets, and subsequently reconstructs the data in a shared space. To create relations between two datasets, the algorithm first identifies mutual nearest neighbors (MNNs). The translation vector is then computed from the resulting list of paired cells (or MNNs) to align the data sets into a shared space. The benefit of this method is that it produces a normalized gene expression matrix, which can be used in downstream analysis and offer an effective correction in the face of compositional variations between batches (Haghverdi, Lun et al. 2018). Scanorama (Hie, Bryson et al. 2019) and batch balanced *k*-nearest neighbors (BBKNN) (Polański, Young et al. 2020) are two other approaches that look for MNNs in reduced-dimension spaces, and use them in a similarity-weighted way to direct batch integration.

Hie et al. (Hie, Bryson et al. 2019) proposed Scanorama which can combine and remove batch effects from heterogeneous scRNA-seq studies by identifying and merging common cell types across all pairs in a dataset. Using a variety of existing tools, Scanorama batch-corrected output can be used for downstream tasks, such as classify cluster-specific marker genes in differential expression analysis. Scanorama outperforms current methods for integrating heterogeneous datasets and it scales to millions of cells, allowing the identification of rare or new cell states through a variety of diseases and biological processes (Hie, Bryson et al. 2019).

Polańsky et al. (Polański, Young et al. 2020) developed BBKNN, a fast graph-based algorithm that removes batch effects through linking analogous cells in different batches. BBKNN is a simple, rapid, and lightweight batch alignment tool, and its output can be directly use for dimensionality reduction. BBKNN’s default approximate neighbor mode scales linearly with the size of datasets and remains consistently faster (by one or two orders of magnitude) when compared to other existing techniques (Polański, Young et al. 2020).

Recently, there has been considerable progress in using DL for batch effect corrections. Residual Neural Networks (ResNets)(He, Zhang et al. 2016) and AEs are two of the most commonly used DL-based batch correction approach in scRNA-seq analysis. ResNets are a form of deep neural network that make a direct connection between the input of a layer (or network) and the outputs, often through an addition operation. Shaham et al. (Shaham, Stanton et al. 2017) suggested a non-linear batch effect correction approach based on a distribution-matching ResNet. Their approach focuses on reducing the Maximum Mean Discrepancy (MMD) between two multivariate replication distributions that were measured in separate batches. Shaham et al. applied their methodology to batch correction of scRNA-seq and mass cytometry datasets, finding that their model can overcome batch effects without altering the biological properties of each sample (Shaham, Stanton et al. 2017).

Li et al. (Li, Wang et al. 2020) presented deep embedding algorithm for single-cell clustering (DESC), an unsupervised DL algorithm for “soft” single-cell clustering which can also remove batch effects. DESC learns a non-linear mapping function from the initial scRNA-seq data space to a low-dimensional feature space using a DNN, iteratively optimizing a clustering objective function. This sequential process transfers each cell to the closest cluster and attempts to account for biological and technical variability across different clusters. Li et al. demonstrated that DESC can eliminate the technical batch effect more accurately than MNN-based methods while better preserving true biological differences within closely related immune cells (Li, Wang et al. 2020).

In a prior study, Shaham (Shaham 2018) proposed batch effect correction through batch-free encoding using an adversarial VAE. Shaham utilizesd the adversarial training to achieve data encoding that corresponded exclusively to a subject’s intrinsic biological state, as well as to enforce accurate reconstruction of the input data. This approach results in maintaining the true biological patterns expressed in the data and minimizing the significant biological information loss (Shaham 2018).

Wang et al. introduced Batch Effect ReMoval Using Deep Autoencoders (BERMUDA) (Wang, Johnson et al. 2019), an unsupervised framework for correcting batch effect in scRNA-seq data across different batches. BERMUDA combines separate batches of scRNA-seq data with completely different cell population compositions and amplifies biological signals by passing information between batches. Most nearest neighbor-based models can manage variations in cell populations between batches when such differences are significant. However, BERMUDA was developed with an emphasis on scRNA-seq data with distinct cell populations in mind, focusing on the similarities between cell clusters rather than significat variations. Altogether, a rapidly expanding number of general DL methods for batch effects correction in biological datasets represent new ways for eliminating batch effects in biological datasets.

##### 3.1.3.3 Dimensionality reduction

Dimensionality reduction is a crucial step in visualizing scRNA-seq data, since typical datasets contain thousands of genes as features (dimensions) (Wang and Gu 2018). The most common dimensionality reduction techniques used for scRNA-seq are principal component analysis (PCA) (Pearson 1901), t-Distributed Stochastic Neighbor embedding (t-SNE) (Van der Maaten and Hinton 2008), diffusion map (Haghverdi, Buettner et al. 2015), Gaussian Process Latent Variable Models (GPLVM) (Titsias and Lawrence 2010, Buettner and Theis 2012), Single-cell Interpretation via Multi-kernel LeaRning (SIMLR) (Wang, Ramazzotti et al. 2017), and Uniform Manifold Approximation and Projection (UMAP) (Becht, McInnes et al. 2019).

In low-dimensional spaces, linear projection methods like PCA traditionally cannot depict the complex structures of single-cell data. On the other hand, nonlinear dimension reduction techniques like t-SNE and UMAP, have been shown to be effective in a variety of applications and are commonly used in single-cell data processing (Ding, Condon et al. 2018). These methods also have some drawbacks, such as lacking robustness to random sampling, inability to capture global structures while concentrating on local data structures, parameter sensitivity, and high computational cost (Zheng and Wang 2019). Several DL techniques for reducing the dimensionality of scRNA-seq data have recently been developed. Here we focus on the ones that are based on VAEs or AEs, which are more commonly used in the field.

Ding et al. (Ding, Condon et al. 2018) proposed a VAE-based model (called scvis) to learn a parametric transformation from a high-dimensional space to a low-dimensional embedding, ultimately learning the estimated posterior distributions of low-dimensional latent variables. Compared to common techniques (e.g. t-SNE), scvis can (i) better obtain the global structure of the data, (ii) provide greater interpretability, and (iii) be more robust to noise or unclear measurements. Ding et al. demonstrated that scvis is a promising tool for studying large-scale and high-resolution single cell populations (Ding, Condon et al. 2018). However, according to Becht et al. (Becht, McInnes et al. 2019), the runtime of scvis is long, particularly for dimensionality reduction, and it appears to be less effective at separating cell populations.

In another work, Wang et al. (Wang and Gu 2018) proposed a method for unsupervised dimensionality reduction and visualization of scRNA-seq using deep VAEs, called VAE for scRNA-seq data (VASC). VASC’s architecture consists of the traditional encoder and decoder network of a VAE, with an addition of a zero-inflated layer that simulates dropout events. In comparison to current methods such as PCA, t-SNE, and Zero Inflated Factor Analysis (ZIFA) (Pierson and Yau 2015), VASC can identify nonlinear patterns present within the data, and has broader compatibility as well as better accuracy, particularly when sample sizes are larger (Wang and Gu 2018).

In 2020, Märtens et al. proposed BasisVAE (Märtens and Yau 2020) as a general-purpose approach for joint dimensionality reduction and clustering of features using a VAE. BasisVAE modified the traditional VAE decoder to incorporate a hierarchical Bayesian clustering prior, demonstrating how collapsed variational inference can identify sparse solutions when over-specifying *K*. (Märtens and Yau 2020).

Peng et al. (Amodio, Van Dijk et al. 2019) proposed an AE-based model that combines gene ontology (GO) and DNNs to achieve a low-dimensional representation of scRNA-seq data. Based on this idea, they proposed two innovative approaches for dimensionality reduction and clustering: an unsupervised technique called “Gene Ontology AutoEncoder” (GOAE) and a supervised technique called “Gene Ontology Neural Network” (GONN) for training their AE model and extracting the latent layer as low dimensional representation. Their findings show that by integrating prior information from GO, neural network clustering and interpretability can be enhanced and that they outperform the state-of-the-art dimensionality reduction approaches for scRNA-seq (Peng, Wang et al. 2019).

In a study by Armaki (Armacki 2018), the dimensionality reduction capabilities of VAE- and AE-based models were evaluated and benchmarked against principal component analysis. They found that the best approach for reducing the dimensionality of single-cell data was using AE-based models, while the more efficient VAEs performed worse in some respects than the linear PCA. One possible hypothesis could be that the prior used for modeling the latent space, which was Gaussian distribution, is not a good fit for single-cell data. A prior more befitting single-cell data (such as negative binomial distribution) could improve the performance of the VAE based model (Armacki 2018).

Finally, there is always the endeavor of optimizing algorithms. As mentioned earlier, developing an efficient DL method for dimensional reduction of data is a necessary next step, since it can potentially improve the quality of lower-dimensional representations.

##### 3.1.3.4 In-Silico Generation and Augmentation

Given limitations on scRNA-seq data availability and the importance of adequate sample sizes, in-silico data generation and augmentation offer a fast, reliable, and cheap solution. Synthetic data augmentation is a standard practice in various ML areas, such as text and image classification (Shorten and Khoshgoftaar 2019). With the advent of DL, traditional data augmentation techniques (such as geometric transformations or noise injection) are now being replaced with deep-learned generative models, primarily VAEs (Kingma and Welling 2013) and GANs (Goodfellow, Pouget-Abadie et al. 2014). In computational genomics, both GAN- and VAE-based models have shown promising results in generating omics data. Here, we focus on the recent methods introduced for generating realistic in-silico scRNA-seq.

Marouf et al. (Marouf, Machart et al. 2020) introduced two GAN-based models for scRNA-seq generation and augmentation called single-cell GAN (scGAN) and conditional scGAN (cscGAN); we collectively refer to these models as scGAN. At the time, scGAN outperformed all other state-of-the-art methods for generating and augmenting scRNA-seq data (Marouf, Machart et al. 2020). The success of scGAN was attributed to the Wasserstein-GAN (Arjovsky, Chintala et al. 2017) that learns the underlying manifold of scRNA-seq data, subsequently producing realistic never-seen-before samples. Marouf et al. showcase the power of scGAN by generating specific cell types that are almost indistinguishable from the real data and augmenting the dataset with the synthetic samples improved the classification of rare cell populations (Marouf, Machart et al. 2020).

In a related work, Heydari et al. (Heydari, Davalos et al. 2021) proposed a VAE-based in-silico scRNA-seq model that aimed at improving Marouf et al.’s training time, stability, and generation quality using only one framework (as opposed to two separate models). Heydari et al. proposed *ACTIVA* (Automated Cell-Type-informed Introspective Variational Autoencoder), which employs a single-stream adversarial VAE conditioned with cell-type information. The cell-type conditioning encourages ACTIVA to learn the distribution of all cell types in the dataset (including rare populations), which allows the model to generate specific cell types on demand. Heydari et al. showed that ACTIVA performs better or comparable to scGAN while training up to 17 times faster due to the design choices. Data generation and augmentation with both ACTIVA and scGAN can enhance scRNA-seq pipelines and analysis, such as benchmarking new algorithms, studying the accuracy of classifiers, and detecting marker genes. Both generative models will facilitate the analysis of smaller datasets, potentially reducing the number of patients and animals necessary in initial studies (Heydari, Davalos et al. 2021).

#### 3.1.4 Downstream Analysis

Following pre-processing, downstream analysis methods are used to derive biological understandings and identify the underlying biological mechanism. For example, cell-type clusters are made up of cells with similar gene expression profiles; minor differences in gene expression between similar cells indicate continuous (differentiation) trajectories; or genes which expression profiles are correlated, signaling co-regulation (Zhang, Cui et al. 2021).

##### 3.1.4.1 Clustering and cell annotation

A significant phase in the scRNA-seq study is to classify cell subpopulations and cluster them into biologically relevant entities (Yang, Liu et al. 2017). The creation of several atlas projects such as Mouse Cell Atlas (Han, Wang et al. 2018), Aging Drosophila Brain Atlas (Davie, Janssens et al. 2018), and Human Cell Atlas (Rozenblatt-Rosen, Stubbington et al. 2017) has been initiated by advances in single-cell clustering. Different clustering approaches have emerged in recent years to help characterize different cell populations within scRNA-seq data (Zheng, Li et al. 2019). Given that DL-based algorithms outperform traditional ML models in clustering tasks when applied to image and text datasets (Guo, Zhu et al. 2018), many have turned to design supervised and unsupervised DL-based clustering techniques for scRNA-seq.

Li et al. presented DESC, an AE-based method for clustering scRNA-seq data with a self-training target distribution that can also denoise and potentially remove batch effects. In experiments conducted by Li et al. (Li, Wang et al. 2020), DESC attained a high clustering accuracy across the tested datasets compared to several existing methods. DESC also showed consistent performance in a variety of scenarios, and did not directly need the batch definition for batch effect correction. Given that Li et al. use a deep AE to reconstruct the input data, the latent space is not regularized with additional properties that could additionally help with clustering (Chen, Wang et al. 2020). As with most DL models, DESC can be trained on CPUs or GPUs.

Chen et al. proposed scAnCluster (Single-Cell Annotation and Clustering), an end-to-end supervised clustering and cell annotation framework that is built upon their previous unsupervised clustering work, namely scDMFK and scziDesk. ScDMFK algorithm (Single-Cell Data Clustering through Multinomial Modeling and Fuzzy K-Means Algorithm) combined deep AEs with statistical modeling. It proposed an adaptive fuzzy k-means algorithm to handle soft clustering while using multinomial distribution to describe the data structure and relying on a neural network to facilitate model parameter estimation (Chen, Wang et al. 2020). More specifically, the AE in scAnCluster learns a low-dimensional representation of the data, and an adaptive fuzzy k-means algorithm with entropy regularization for soft clustering. On the other hand, ScziDesk aimed to learn a “cluster-friendly” lower-dimensional representation of the data. Many existing DL-based clustering algorithms for scRNA-seq do not consider distance and affinity constraints between similar cells. This prohibits such models to learn cluster friendly lower-dimensional representations of the data (Chen, Wang et al. 2020). For scAnCluster, Chen et al. use the available cell marker information to construct a new DL model that integrates single-cell clustering and annotation. ScAnCluster can do both intra-dataset and inter-dataset cell clustering and annotation, and it also reveals a clear discriminatory effect in the detection of new cell types not found in the reference (Tran, Ang et al. 2020).

Tian et al. created the Single-Cell model-based Deep embedded Clustering (scDeepCluster) technique, which uses a nonlinear approach to combine DCA modeling and the DESC clustering algorithm. Their approach sought to improve clustering while reducing dimensions directly. ScDeepCluster outperformed state-of-the-art approaches on a variety of clustering efficiency metrics, showing runtime increasing linearly with sample size. In contrast to similar methods, scDeepCluster requires less memory and is scalable to larger datasets (Tian, Wan et al. 2019). On the other hand, ScDeepCluster lacks the pairwise distance of associated cells and ignores the affinity constrains of similar cells. ScDeepCluster does not pre-select informative genes as input data, making the model more computationally efficient, but decreasing the clustering accuracy (Chen, Wang et al. 2020).

Peng et al. (Peng, Wang et al. 2019) developed a strategy for optimizing cell clustering based on global transcriptome profiles of genes. They used a combination of DNN and GO to reduce the dimensions of scRNA-seq data and improve clustering. Their supervised approach was based on a conventional neural network, while the unsupervised model utilized an AE. Their model consisted primarily of two main components: the choosing of important GO factors and the combination of GO terms with the DNN-based model (Peng, Wang et al. 2019). Single-Cell Variational Auto-Encoders (scVAE) (Grønbech, Vording et al. 2020) is another VAE-based model for downstream analysis scRNA-seq data. However, compared to other models, scVAE has the advantage of not requiring many of the traditional pre-processing steps since it uses the raw count data as input. ScVAE can accurately predict expected gene expression levels and a latent representation for each cell and is flexible to use the known scRNA-seq count distributions (such as Poisson or Negative Binomial) as its model assumption (Grønbech, Vording et al. 2020).

##### 3.1.4.2 Cell-Cell communication analysis

In recent years, scRNA-seq has become a powerful tool for analyzing cell-cell communication in tissues. Intercellular communication controlled by ligand-receptor complexes is essential for coordinating a wide range of biological processes, including development, differentiation, and inflammation. Several algorithms have been proposed to carry out these analyses. These algorithms begin with a database of interacting molecular partners (such as ligand and receptor pairs) and predict a list of possible signaling pathways between types of cells based on their expression patterns. Although the findings of these studies may provide useful insights into the mechanisms of tissues, they can be difficult to visualize and analyze using current algorithms (Tian, Wan et al. 2019, Almet, Cang et al. 2021). DL-based algorithms in this space have not yet been fully developed. However, we expect new DL methods to emerge in the near future, given the importance of cell-cell communication analysis.

##### 3.1.4.3 RNA velocity

RNA velocity has created new ways to research cellular differentiation in scRNA-seq studies. RNA velocity represents the rate of change in gene expression for a single gene at a given time point depending on the spliced ratio to unspliced mRNA. In other words, RNA velocity is a vector that forecasts the possible future of individual cells on a timescale of hours, which prove new insights into cellular differentiation that have challenged conventional and long-standing views. Existing experimental models for tracking cell fate and reconstructing cell lineages, such as genetic methods or time-lapse imaging have limited power as they do not show the trajectory of differentiation or reveal the molecular identity of intermediate states. But recently, with the advent of scRNA-seq, a variety of algorithms such as VeloViz (Atta and Fan 2021) or scVelo (Bergen, Lange et al. 2020) can visualize the velocity estimates in low dimensions. Although such models show promising results (particularly in well-characterized systems), we are far from a complete understanding of cellular differentiation and cell fate decisions. By this means, we foresee the development of more integrative models, such as DL-based approaches, to resolve long-standing concerns regarding cell fate choices and lineage specification using RNA velocity.

### 3.2 Deep Learning in Single-Cell Genomics

Traditional sequencing methods are limited to measuring the average signal in a group of cells, potentially masking heterogeneity and rare populations (Xiong, Xu et al. 2019). On the other hand, scRNA-seq technologies provide a tremendous advantage for investigating cellular heterogeneity and recognizing new molecular features correlated with clinical outcomes, resulting in the transformation of many biological research domains. Similar to scRNA-seq, single-cell (SC) genomics is being used in many areas, such as predicting the sequence specificity of DNA- and RNA-binding proteins, enhancer and cis-regulatory regions, methylation status, gene expression, control splicing, and searching for associations between genotype and phenotype. However, SC genomics data are often too large and complex to be analyzed only through visual investigation of pairwise correlations. As a result, a growing number of studies have leveraged DL techniques to process and analyze these large datasets. In addition to the scalability of DL algorithms, another advantage of DL techniques is the learned representations from raw input data, which are beneficial in specific SC genomics and epigenomics applications. These application include cell-type identification, DNA methylation, chromatin accessibility, TF-gene relationship prediction, and histone modifications. In the following sections, we review some applications of DL models for analyzing scDNA-seq data (summarized in Fig. 4).

**Figure 3:**
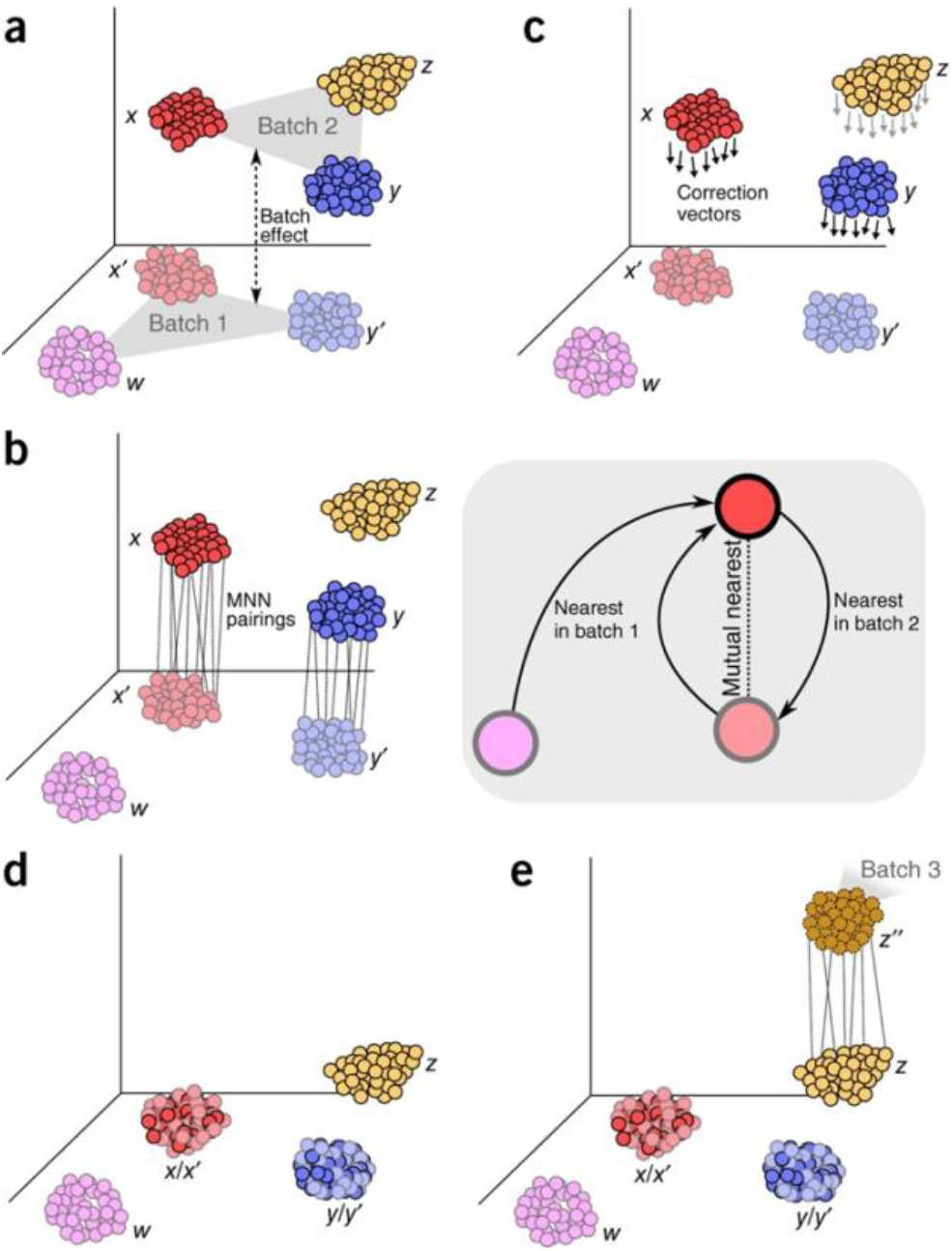
Batch-effect correction via MNN. (a) Batch 1 and batch 2 in high dimensions, with a batch effect variation that is almost orthogonal. (b) By identifying MNN pairs of cells, the algorithm recognizes matching cell types (gray box). (c) Between the MNN pairs, batch-correction vectors are measured. (d) Batch 1 is considered the reference, and batch 2 is combined into it by subtracting correction vectors. (e) The integrated data is used as a reference, and the process is repeated with each new batch of data. This figure has been reused with permission from authors (Haghverdi, Lun et al. 2018).

**Figure 4.**
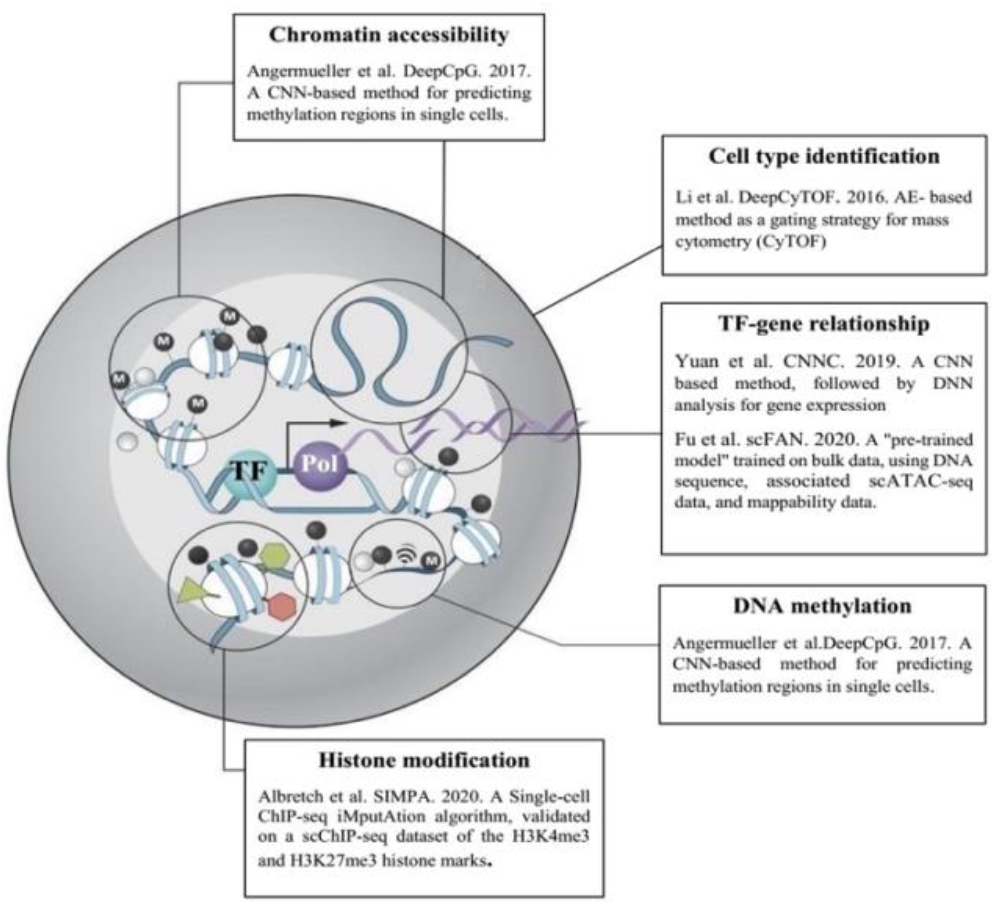
Overview of algorithms that are used in different parts of single-cell genomics analysis.

#### 3.2.1 Cell type identification in CyTOF

An essential and challenging task in genomics research is to accurately identify and cluster individual cells into distinct groups of cell types. Li et al. (Li, Shaham et al. 2017) describe AE methods (stacked AE and multi-AE) as a gating strategy for mass cytometry (CyTOF). CyTOF is a recent technology for high-dimensional multiparameter SC analysis. They introduced DeepCyTOF as a standardization procedure focused on a multi-AE neural network. DeepCyTOF focuses on domain adaptation principles and is a generalization of previous work that helps users in calibrating between a source domain distribution (reference sample) and several target domain distributions (target samples) in a supervised manner. Moreover, DeepCyTOF requires labelled cells from only a single sample. DeepCyTOF was applied to two CyTOF datasets produced from primary immune blood cells: (a) cases with a history of West Nile virus (WNV) infection and (b) normal cases of various ages. They manually gated a single baseline reference sample in each of these datasets to automatically gate the remaining uncalibrated samples.

Li et al. revealed that DeepCyTOF’s cell classification was very consistent with classifications done by individual manual gatings, with over 99% concordance. Additionally, they used a stacked AE (which is one of DeepCyTOF’s key components) to tackle the semi-automated gating challenge of the FlowCAP-I competition. Li et al. found that their model outperformed other existing gating approaches benchmarked on the fourth challenge of the competition. Overall, stacked AEs combined with a domain adaptation technique suggest promising results for CyTOF semi-automated gating and flow cytometry data, requiring manual gating of one reference sample to precisely gate the remaining samples.

#### 3.2.2 DNA methylation

Recent technological advances have made it possible to assay DNA methylation at the single-cell resolution. Angermueller et al. propose DeepCpG (Angermueller, Lee et al. 2017), a CNN-based computational method for predicting methylation regions. DeepCpG is made up of three modules: a DNA module that extracts features from the DNA sequence, a CpG module that extracts features from the CpG neighborhood of all cells, and a multi-task joint module that integrates evidence from both modules for predicting the methylation regions of target CpG sites for different cells. The trained DeepCpG model can be used in various downstream studies, such as inferring low-coverage methylation profiles for groups of cells and recognizing DNA sequence motifs linked to methylation states and cell-to-cell heterogeneity. Angermueller et al. apply their model to both mouse and human cells, which achieves significantly better predictions than previous techniques (Angermueller, Lee et al. 2017). An overview of the DeepCpG model is shown in (Fig. 5).

**Figure 5.**
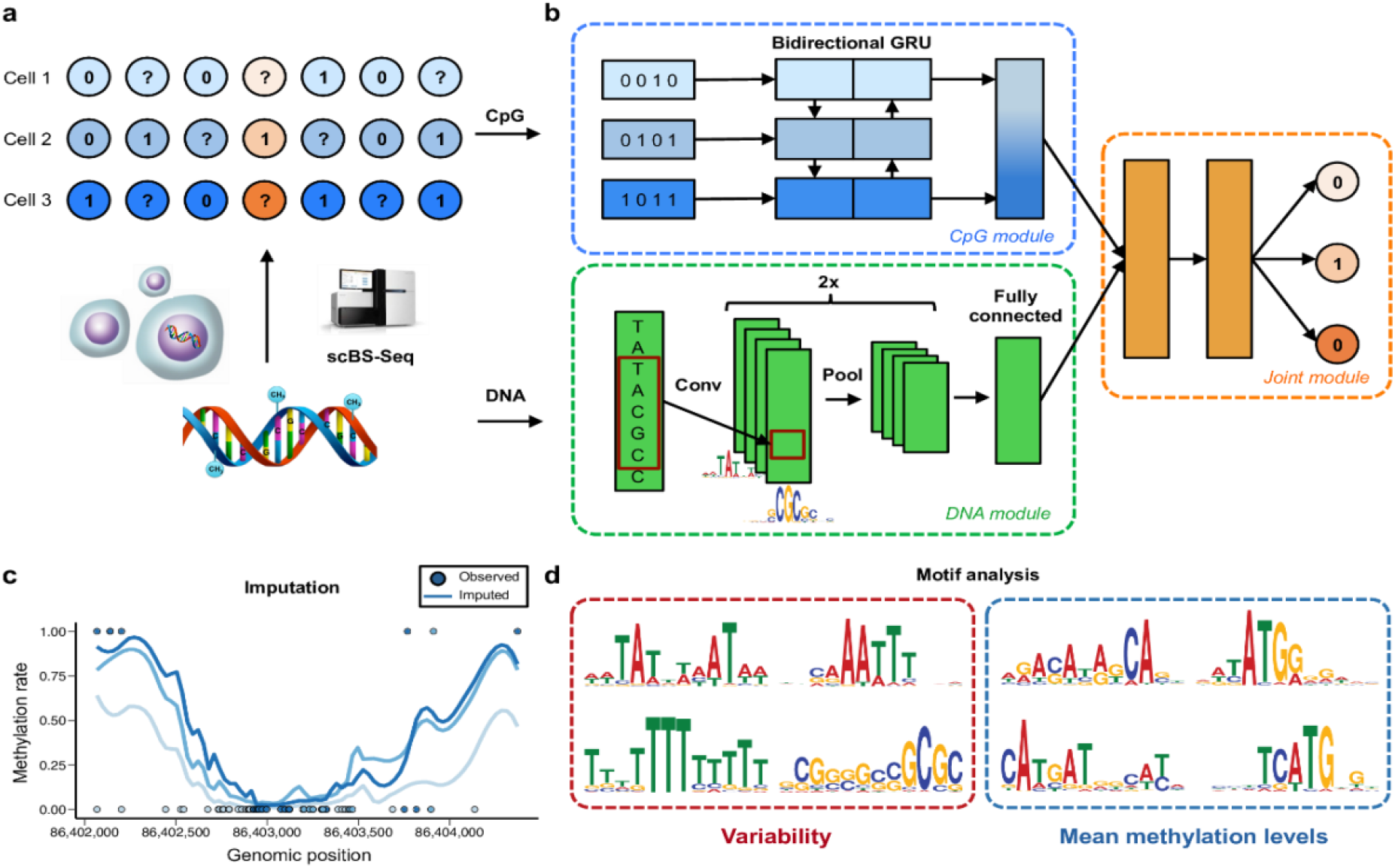
An overview of DeepCpG (taken from Angermueller et al.) (a) scBS-seq (Smallwood et al.) scRRBS-seq (Farlik et al.) provide sparse single-cell CpG profiles, which are then pre-processed as binary values (methylated CpG sites are marked by ones, unmethylated CpG sites are denoted by zeros, and CpG sites with unknown methylation state [missing data] are represented by question marks) (b) Two convolutional and pooling layers are used to detect predictive motifs from the local sequence context, and one fully connected layer is used to model motif interactions in the DNA model. Applying a bidirectional gated recurrent network (GRU), the CpG model scans the CpG neighborhood of numerous cells (rows in b), providing compressed information in a vector of constant size. To forecast methylation states in all cells, the Joint model learns connections between higher-level properties generated from the DNA- and CpG models (c, d) The DeepCpG model can be utilized for a variety of downstream tasks after training, such as genome-wide imputation of missing CpG sites (c) and the finding of DNA sequence motifs linked to DNA methylation levels or cell-to-cell variability (d).

Interestingly, DeepCpG can be used for differentiating human induced pluripotent stem cells in parallel with transcriptome sequencing to specify splicing variation (exon skipping) and its determinants. Linker et al. (Linker, Urban et al. 2019) presented that variation in SC splicing can be precisely predicted based on local sequence composition and genomic features. DeepCpG, which is used for DNA methylation profiles, imputes unobserved methylation regions of individual CpG sites. The cell-type-specific models were made using CpG and genomic information according to DeepCpG’s setup of a joint model. Finally, during cell differentiation, Linker et al. identified and characterized associations between DNA methylation and splicing changes, which led to indicating novel insights into alternative splicing at the SC level.

Another method for studying chromatin accessibility at single-cell resolution, called Assay of Transposase Accessible Chromatin sequencing (scATAC-seq), has recently gained considerable popularity. In scATAC-seq, mutation-induced hyperactive Tn5 transposase tags and fragments regions in open chromatin sites in the DNA sequence, which is later sequenced using paired-end Next Generation Sequencing (NGS) technologies (Yan, Powell et al. 2020). The pre-processing steps of scATAC-seq data analysis are often analogous to scRNA-seq pipelines. That is, the same tools are often used in both data modalities, although scRNA-seq tools are often not optimized for the particular properties of scATAC-seq data. ScATAC-seq has low coverage, and the data analysis is highly sensitive to non-biological confounding factors. The data is pre-processed and assembled into a feature-per-cell matrix, where common choices for “feature” are fixed-size genomic bins and signal peaks at biological events. This matrix displays particular numerical properties which entail computational challenges: it is extremely high-dimensional, sparse, and near-binary in nature (presence/absence of signal). Several packages have been recently developed specifically for scATAC-seq data, with all of them having some major limitations (Cao, Fu et al. 2021). The tools for processing scATAC-seq are diverse and nescient, and thus, there is no consensus on the best practices scATAC-seq data analysis. Further development of scATAC-centric computational tools and benchmark studies are much needed. ScATAC-seq analyses can help to elucidate cell types and differentially accessible regions across cells. Moreover, it can decipher regulatory networks of cis-acting elements like promoters and enhancers, and trans-acting elements like transcription factors (TFs), and infer gene activity (Baek and Lee 2020). ScATAC-seq data could also be integrated with RNA-seq and other omics data. However, most current software only integrates the derived gene activity matrix with expression data, and important information from whole-genome chromatin accessibility is lost.

Currently, there are a variety of DL models for bulk ATAC-seq data, such as LanceOtron’s CNN for peak calling (Hentges, Sergeant et al. 2021) and CoRE-ATAC for functional classification of cis-regulatory elements (Thibodeau, Khetan et al. 2020). Thibodeau et al. demonstrated CoRE-ATAC’s transferable capability on cell clusters inferred from single nuclei ATAC-seq data with a small decrease in model prediction accuracy (mean micro-average precision of 0.80 from bulk vs. 0.69 in single-cell clusters).

One common way to reduce the dimensionality of scRNA-data is to identify the most variable genes (e.g. with PCA), since they carry the most biologically relevant information. However, scATAC-seq data is binary, and therefore prohibiting the identification of variable peaks for dimensionality reduction. Instead, dimensionality reduction of scATAC data is done through Latent Semantic Indexing (LSI), a technique used for natural language processing. Although this approach is scalable to large number of cells and features, it may fail to capture the complex reliance of peaks, since LSI is a linear method.

Single-Cell ATAC-seq analysis via Latent feature Extraction (SCALE) (Xiong, Xu et al. 2019) combines a deep generative framework with a probabilistic Gaussian Mixture Model (GMM) as a prior over the latent variables, in order to learn a nonlinear latent space of scATAC-seq features. Given the nature of scATAC-seq, GMM is a suitable distribution for modeling the high-dimensional, sparse multimodal scATAC-seq data. SCALE can also be used for denoising and imputing missing data, recovering signals from missing peaks. In the benchmarking study by Xiong et al., they demonstrated that SCALE outperformed conventional non-DL scATAC-seq tools for dimensionality reduction, such as PCA and LSI. Furthermore, they show that SCALE is scalable to large datasets (on the order of 80,000 single cells). While SCALE succeeds at learning nonlinear cell representations with higher accuracy, it assumes that the read depth is constant across cells and ignores potential batch effects. These pitfalls motivated the development of scalable and accurate invariant representation learning scheme (SAILER) (Cao, Fu et al. 2021).

SAILER is a deep generative model inspired by VAEs which also learns a low-dimensional latent representation of each cell. For SAILER, the authors aimed to design an invariant representation learning scheme, where they discard the learned component associated with confounding factors from various technical sources. SAILER captures nonlinear dependencies among peaks, faithfully separating biologically relevant information from technical noise, in a manner that is easily scalable to millions of cells (when using GPUs). Similar to SAILER, SCALE also offers a unified strategy for scATAC-seq denoising, clustering, and imputation. However, in multi-sample scATAC-seq integration, SAILER can eliminate batch effects and properly recreate a chromatin accessibility landscape free of confusing variables, regardless of sequencing depths or batch effects.

#### 3.2.3 Transcription Factor (TF)-gene relationship prediction

To unravel gene regulatory mechanisms and differentiate heterogeneous cells, understanding the genome-wide binding TF profile is crucial. Researchers have developed several methods that use expression data for infer gene-gene interactions such inferring coexpression, understanding functional assignments and reconstructing pathways (Yuan and Bar-Joseph 2019). However, each task in infering gene-gene relationships is typically done using different techniques. DL-based methods, on the other hand, are capable of learning multiple tasks jointly, which is very advantageous in inferring relationships between genes. Yuan et al. (Yuan and Bar-Joseph 2019) introduced Convolutional Neural Network for Co-Expression (CNNC), a new encoding method for gene expression data based on CNNs. CNNC is a general computational technique for supervised gene relationship inference that builds on previous approaches in various tasks, including predicting TF targets and recognizing genes related to disease in order to infer cause and effect. The key idea behind CNNC is to turn data into co-occurrence histograms (as images), and then analyzing the histograms using CNNs. More specifically, Yuan et al. generated a histogram for each pair of genes and utilized CNNs to infer relationships among the various levels of expression encoded in the image. CNNC is adaptable and can easily be expanded to integrate other types of genomics data as well, resulting in additional performance gains. CNNC goes beyond previous approaches for predicting TF-gene and protein-protein interactions and predicting the pathway of a regulator-target gene pair. CNNC may also be used to draw causality inferences, functional assignments (such as biological processes and diseases), and as part of algorithms that recreate known pathways (Yuan and Bar-Joseph 2019).

In another work, Fu et al. (Fu, Zhang et al. 2020) presented Single Cell Factor Analysis Network (scFAN), a DL model for determining genome-wide TF binding profiles in individual cells. The scFAN pipeline consists of a “pre-trained model” trained on bulk data and then used to predict TF binding at the cellular level using DNA sequence, aggregated associated scATAC-seq data, and mappability data. ScFAN can help overcome the basic sparsity and noise constraints of scATAC-seq data. This model provides a valuable method for predicting TF profiles through individual cells and could be applied to analyze SC epigenomics and determine cell types. Fu et al. presented scFAN’s ability to identify cell types by analyzing sequence motifs enriched within predicted binding peaks and studying the effectiveness of predicted TF peaks. They suggested a novel metric called “TF activity score” to classify each cell and demonstrated that the activity scores could accurately capture cell identity. Generally, scFAN is capable of connecting open chromatin states with transcript factor binding activity in individual cells, which is beneficial for a deeper understanding of regulatory and cellular dynamics (Fu, Zhang et al. 2020).

#### 3.2.4 Histone modification

Given the effects of protein-DNA interactions between histone marks and TF on the regulation of crucial cellular processes (including the organization of chromatin structures and gene expression), the identification of such interactions is highly significant in biomedical science. Chromatin immunoprecipitation followed by sequencing (ChIP-seq) is a widely used technique for mapping TFs, histone changes, and other protein-DNA interactions for genome-wide mapping (Furey 2012). ChIP-seq data is very sparse, therefore generally requiring imputation for more accurate analysis. Albretch et al. introduced Single-cell ChIP-seq iMPutAtion (SIMPA) (Albrecht, Andreani et al. 2021), an imputation algorithm that was tested on a single-cell ChIP-seq (scChIP-seq) dataset of the H3K4me3 and H3K27me3 histone marks in B-cells and T-cells. Unlike most SC imputation approaches, SIMPA integrates the sparse input of one single cell with a series of 2,251 ENCODE ChIP-seq experiments to extract predictive information from bulk ChIP-seq data. SIMPA’s goal is to identify statistical patterns that bind protein-DNA interacting sites through specific SC regions of target-specific ENCODE data for different cell types, as well as the presence or absence of potential sites for a single cell. Once the patterns are identified, SIMPA’s DL models uses these patterns to make precise predictions. As a new approach in sc-seq, SIMPA’s imputation strategy augmented sparse scChIP-seq data, leading to improved cell-type clustering and the detection of pathways that are specific to each cell type.

SC genomics analysis through DL is a promising and emerging field with an incredible potential to advance our knowledge of fundamental biological matters. In this respect, DL can provide us with a better understanding of nature, and the intricacies of DNA structure and epigenomics effects on human diseases for both therapeutic and diagnostic purposes. Due to intrinsic challenges in SC genomics, such as sparsity, systematic noise, and higher-dimensionality of biological systems, developing new DL models are paramount to further advancing the SC genomics field.

### 3.3 Deep Learning in Spatial Transcriptomics

Since being named the method of the year (Marx 2021), spatial transcriptomics (ST) is becoming the natural extension of scRNA-seq, unbiasedly profiling transcriptome-wide gene expression. By not requiring tissue dissociation, spatial transcriptomic retain spatial information (adding a spatial component to conventional RNA-seq technologies). ST have the potential of revolutionizing the field by bridging the gap between the deep characterization of cellular states and the cellular diversity that constitutes tissue organization. Spatially resolved transcriptomics can provide the genetic profiles of cells while containing information about the positional distribution of the sequenced cells, enhancing our understanding of cell interactions, organ function and pathology. However, the high-throughput spatial characterization of complex tissues remains a challenge. Broadly, spatially-resolved transcriptomics techniques can be divided into two categories: (i) the cyclic RNA imaging techniques that achieve single-cell resolution and (ii) array-based spatially resolved RNA-seq techniques, such as Visium Spatial Transcriptomics (Ståhl, Salmén et al. 2016), Slide-sequencing (Rodriques, Stickels et al. 2019), or high-definition spatial transcriptomics (HDST) (Vickovic, Eraslan et al. 2019). Though both subgroups can provide spatial information on a single-cell level, the cyclic RNA methods are limited on the number of genes that they can multiplex. On the other hand, the array-based spatially resolved RNA-seq techniques achieve high-throughput data by capturing mRNA across thin tissue sections using a grid of microarrays or bead-arrays, relying on simple molecular biology and histology protocols. However, since array-based mRNA capture does not match cellular boundaries, spatial RNA-seq measurements are a combination of multiple cell-type gene expressions that can correspond either to multiple cells (Visium) or fractions of multiple cells (depending on the spatial resolution of each method).

To obtain a comprehensive characterization of underlying tissue, there is the need for computational methods that can produce coupled single-cell and spatially resolved transcriptomics strategy, mapping cellular profiles into a spatial context. Nonetheless, there are some techniques for retrieving relevant biological information from ST data. As remarked by Lähnemann et al. (Lähnemann, Köster et al. 2020), detecting spatial gene expression patterns is one of the most pressing challenges in single-cell omics data science. Identifying such patterns can provide valuable insight on the spatial distribution of cell populations, pointing out gene marker candidates and potentially leading to identification of new rare cell subpopulations. Moreover, ST not only puts gene expression into a spatial context but also facilitates the integration of tissue-image information with gene expression information. Such data integration will enable researchers to utilize image processing techniques to investigate the morphological information, gaining more intuition in order to obtain more refined inferences, predictions, or cellular profiles. After addressing the mapping issues from gene expression to the spatial coordinates, further computational tools will be required to study cell-cell interactions within tissues and to model transcriptional relations between cell types (Pham, Tan et al. 2020). Given the recency of ST, ML- and DL-based models for studying this type of data are rare and not fully developed. Given the complexity of the ST space, however, we predict that DL models will be the predominant method of choice for ST data integration and analysis, with perhaps many new models borrowing ideas and designs from the existing body of work in computer vision.

### 3.4 Integrating scRNA-seq and Spatial Transcriptomics (Spot Deconvolution)

Given that ST approaches normally detect mRNA expression from a mixture of cells, and that they do not distribute the sequenced samples to match cellular boundaries, it is imperative to integrate ST data with scRNA-seq data to obtain a comprehensive map. Using scRNAs-seq as a reference, this integration aims to infer which cells belong to which gene expression counts detected at the various location across the tissue. Thus far, traditional ML and statistical models have shown promising results in this space. For example, SPOTlight (Elosua-Bayes, Nieto et al. 2021) is a model for deconvoluting spatial transcriptomics spots with single-cell transcriptomes using seeded NMF regression. SPOTlight starts by obtaining cell-type-specific profiles that are representatives of the gene expressions associated with the different cells. This process is further refined by a proper initialization of the method that uses unique marker genes of specific cell types. Next, the method uses Non-Negative Least Squares to deconvolute the captured expression of each spot (location). Despite SPOTlight’s accuracy and computational efficiency, SPOTlight lacks the flexibility of integrating datasets from different batches or sequencing technologies. In SPOTlight, several aspects of the underlying biological and technological variance in the data are not addressed nor accounted for, which limits its application. Furthermore, technical procedures to obtain RNA-seq data are notoriously delicate and subjected to notable sources of variation.

Methods like Seurat 3 (Stuart, Butler et al. 2019) attempt to account for the intrinsic technical variability of RNA-seq procedures thorough developing an “anchor”-based method for integrating datasets. However, it is important to remember that there are biological variabilities in the number of cells across positions or the amount of mRNA expressed by each cell or cell type. Therefore, there is an unmet need for techniques that can integrate datasets while properly accounting for the various sources of variability. Several studies have proposed statistical frameworks as viable candidates to account for the variability when integrating different datasets. For example, Stereoscope (Andersson, Bergenstråhle et al. 2020) build their model on a statistical framework. They model gene expressions counts as occurrences under a negative binomial distribution. Stereoscope follows previous approaches of obtaining a gene expression profile for each cell type. That is, Andersson et al. follow a two-step approach: First, they estimate the parameter of the negative binomial distribution for all genes within each cell type. Similar parameters for a distribution of the RNA-seq and spatial expression mixture are then formed by a linear combination of the single-cell parameters. The next step is to search a set of weights that can best fit the spatial data (Andersson, Bergenstråhle et al. 2020). These computed weights reflect the contribution of each cell type to the gene expression counts found in each location, thus explaining the abundance of each cell type across the spots.

Following the previous methodology, cell2location (Kleshchevnikov, Shmatko et al. 2020) builds on a Bayesian framework and models gene expression counts as a negative binomial distribution. This approach enables controlling the sources of variability, which is crucial when working with data from different technologies. Cell2location integrates scRNA-seq information into the spatial model using a similar statistical framework as Stereoscope: In addition to modeling the gene-specific unobserved rate (mean) as a weighted sum of the cell signature gene expression, cell2location also adds various parameters to provide the model with prior information regarding technology sensitivity. That is, to further improve integration between different technologies, Kleshchevnikov et al. allow for four hyperparameters that will scale the weighted sum of cell contributions. Kleshchevnikov et al. showed that cell2location results in a more comprehensive integration of omics data while being more computationally efficient than competing models (such as Stereoscope).

### 3.5 Deep Learning in the Integration of Single-Cell Multimodal Omics Data

Single-cell sequencing (scSeq) was chosen as Method of the Year in 2013 due to its ability to sequence DNA and RNA in individual cells (Teichmann and Efremova 2020). ScSeq allows for gene expression measurements at an unprecedented single-cell resolution, which can provide a comprehensive view of the genome, transcriptome, or epigenome. In addition, recent technological advances now allow for multimodal omics measurements from the same experiment. In order to gain an accurate and comprehensive view of the cellular composition of the control (normal) and disease groups, it is essential to integrate all omics data for one sample simultaneously (Wani and Raza 2019). The various omics technologies can assess various modalities in one experiment (i.e. conduct multimodal studies) or integrate diverse omics datasets from multiple experiments.

Given the tremendous potentials of these approaches, single-cell multimodal omics was named the 2019 Method of the Year (Teichmann and Efremova 2020). Omics integration holds the promise of linking even small datasets across orthogonal biochemical domains, amplifying biologically significant signals in the process (Grapov, Fahrmann et al. 2018). By analyzing multi-omics data, researchers can produce novel hypotheses or design mathematical algorithms for prediction tasks, such as drug sensitivity and efficacy, gene dependence prediction, and patient stratification. Hao et al (Hao, Hao et al. 2021) proposed a non-DL based method, called the weighted-nearest neighbor method which has shown promising results. WNN is an unsupervised framework for defining cellular identity by leveraging multiple data types for creating a multimodal reference atlas. Hao et al. apply their technique to a dataset of human PBMCs that includes paired transcriptomes and measurements of 228 surface proteins, forming a multimodal immune system atlas. They evaluate multimodal datasets from single cells, including paired measurements of RNA and chromatin state, and extend beyond the transcriptome to define cellular identity in a coherent and multimodal manner (Hao, Hao et al. 2021). Although traditional ML models such as WNN have shown promising results, multi-omics data pose unique challenges for holistic integration, in addition to the other traditional difficulties such as batch effects from multiple sources. Multi-omics data reflect molecular phenotypes at different molecular systems, and thus each omics dataset could follow a different and specific distribution. To overcome these hurdles, sophisticated statistical and computational strategies are required. Among the various proposed algorithms thus far, only DL-based algorithms provide the computational versatility necessary to effectively model and incorporate virtually any form of omics data in an unsupervised or supervised manner (Grapov, Fahrmann et al. 2018).

Most DL-based algorithms in this area aim to simultaneously calculate several modalities in a single experiment. For example, Zuo et al. (Zuo and Chen 2020) introduced a single-cell multimodal VAE model (scMVAE) for profiling both transcriptomic and chromatin accessibility information in the same individual cells. Given the scRNA-seq and scATAC-seq of the same individual cells, scMVAE’s uses three joint-learning strategies to learn a non-linear joint embedding that can be used for various downstream tasks (e.g. clustering). This joint learning distinguishes scMVAE from other VAE-based models (such as scVI) which process individual omics data separately. Zuo et al. note that scMVAE’s feature embeddings are more distinct than the scVI’s for each omics data, indicating that the joint learning representation of multi-omics data will produce a more robust and more valuable representation (Zuo and Chen 2020).

Currently, there are only a few studies that have used DL for data integration, but their success thus far calls for additional investigation of DL models in this domain. Despite the significant advances made with single-cell multimodal omics technologies, several obstacles remain: First, these techniques are prohibitively expensive when used on a large scale to analyze complex heterogeneous samples and distinguish rare cell types within a tissue. On the other hand, data sparsity is a significant limitation of high-throughput single-cell multimodal omics assays. Furthermore, existing methods cover only a small portion of the epigenome and transcriptome of individual cells, making it challenging to separate technical noise from cell-to-cell variability. While future modification of these approaches will eventually close the gap, fundamentally new algorithms or strategies may be needed to resolve this constraint completely (Zhu, Preissl et al. 2020). Amodio et al. (Amodio and Krishnaswamy 2018) propose Manifold-Aligning GAN (MAGAN), which is a GAN-based model which aligns two manifolds coming from different domains with the assumption that different measurements for the same underlying system contain complementary information. They show MAGAN’s potential in the problem of single-cell data integration (CyTOF and scRNA-seq data) and demonstrate the generalization potential of this method in the integration of other data types. However, the performance of MAGAN decreases in the absence of correspondence complementary information among samples (Liu, Huang et al. 2019). Cao et al. introduced UnionCom for unsupervised topological alignment of single-cell omics integration without a need for correspondence information among cells or among features, which can be very useful in data integration. However, UnionCom is not scalable to large datasets that are on the order of millions of cells (Cao, Bai et al. 2020). Other models such as SMILE (Single-cell Mutual Information Learning) (Xu, Das et al. 2021) also allow for unmatched feature types. SMILE is a deep clustering algorithm for different tissues and modalities, even when the feature types are unmatched. SMILE can remove batch effects and learn a discriminative representation for data integration using a cell-pairing maximization algorithm (Xu, Das et al. 2021).

Single-Cell Data Integration via Matching (SCIM) (Stark, Ficek et al. 2020) is another deep generative approach that constructs a technology-invariant latent space to recover cell correspondences among datasets, even with unpaired feature sets. The architecture is a modified auto-encoder with an integrated discriminator network, similar to the one in GANs, allowing the network to be trained in an adversarial manner. Multi-modal datasets are integrated by pairing cells across technologies using a bipartite matching scheme that operates on the low-dimensional latent representations (Stark, Ficek et al. 2020). Another data integration model is inteGrative anaLysis of mUlti-omics at single-cEll Resolution (GLUER) (Peng, Chen et al. 2021), which employs three computational approaches: a nonnegative matrix factorization (NMF), mutual nearest neighbor, and a DNN to integrate multi-omics data. The NMF stage helps to identify shared signals across data sets of different modalities, resulting in “factor loading matrices” (FLM) for each modality. The FLM from one data modality is defined as the reference matrix while the other FLM is used as query matrices, which are used to calculate putative cell pairs. These cell pairs are then used in a DNN which tries to learn a mapping between the query FLMs and the reference FLMs, resulting in co-embedded datasets (Peng, Chen et al. 2021).

Some studies have formulated the integration problem as a transfer-learning task. For example, Lin et al. (Lin, Wu et al. 2021) pose the integration problem as a transfer learning question, where the model is co-trained on labeled RNA and unlabeled ATAC data. Their model, scJoint, integrates atlas-scale collections of scRNA-seq and scATAC-seq data, using a neural network framework (Lin, Wu et al. 2021). Their approach uses scATAC-seq to gain a complementary layer of information at the single-cell resolution which is then added to the gene expression data from scRNA-seq. However, scJoint requires that both input matrices share the same dimensions, and scATAC-seq is first converted to gene activity scores, where a single encoder can share the weights for both RNA and ATAC data (Lin, Wu et al. 2021).

## 4. Conclusion

Single-cell (SC) omics technologies generate large datasets that describe the genomic, transcriptomic, or epigenomic profiles of many individual cells in parallel. Integrative methods have recently opened a new way for delineating heterogeneous mechanistic landscapes and cell-cell interactions in single-cell multi-omics. Given the challenges in inferring biological information and constructing predictive models for such datasets, there has been a surge in the use of DL-based models for SC omics. So far, deep learned models have shown promising results, demonstrating the ability to process and learn from massive quantities of high-dimensional representations of single-cell and multi-omics data. However, a thorough understanding of the underlying models is needed to evaluate the optimal pipeline for analyzing single-cell datasets to understand cellular identity and function better. We believe that the sc-seq space will pose unique challenges in representation and DL, particularly in multi-omics applications.

The value and potentials of DL in SC omics have been further highlighted during the current Coronavirus (COVID-19) pandemic. Many scientists have utilized deep-learned models in SC analysis for diagnosing and predicting the severity of COVID-19 (Jeong, Jia et al. 2021), predicting future mutations (Saha, Ghosh et al. 2021) and testing drug combinations for treating COVID-19 (Jin, Stokes et al. 2021). Researchers have also used DL in studying the immune dysfunction in COVID-19 patients, particularly PBMCs and lung tissues, which has allowed scientists to better characterize COVID-19 (Han and Zhang 2021). Overall, the advances of the DL techniques in SC have provided researchers with a unique ability to develop new therapeutic interventions, contributing to a global coordinated effort to win the fight against diseases, such as COVID - 19.

We anticipate an increase in the use of DL techniques for addressing the current challenges mentioned in the SC field. Furthermore, we believe that the biological generalizability and interpretability of deep-learned models in understanding complex pathological phenotypes such as cancer, drug resistance, and neurobiology would be of great interest and importance to the omics field.

## References

Albrecht, S., T. Andreani, M. A. Andrade-Navarro and J.-F. Fontaine (2021). “Interpretable machine learning models for single-cell ChIP-seq imputation.” BioRxiv: 2019.2012. 2020.883983.

Alipanahi, B., A. Delong, M. T. Weirauch and B. J. Frey (2015). “Predicting the sequence specificities of DNA-and RNA-binding proteins by deep learning.” Nature biotechnology 33(8): 831–838.

Almet, A. A., Z. Cang, S. Jin and Q. Nie (2021). “The landscape of cell-cell communication through single-cell transcriptomics.” Current opinion in systems biology.

Amodio, M. and S. Krishnaswamy (2018). MAGAN: Aligning biological manifolds. International Conference on Machine Learning, PMLR.

Amodio, M., D. Van Dijk, K. Srinivasan, W. S. Chen, H. Mohsen, K. R. Moon, A. Campbell, Y. Zhao, X. Wang and M. Venkataswamy (2019). “Exploring single-cell data with deep multitasking neural networks.” Nature methods 16(11): 1139–1145.

Anchang, B., T. D. Hart, S. C. Bendall, P. Qiu, Z. Bjornson, M. Linderman, G. P. Nolan and S. K. Plevritis (2016). “Visualization and cellular hierarchy inference of single-cell data using SPADE.” Nature protocols 11(7): 1264–1279.

Andersson, A., J. Bergenstråhle, M. Asp, L. Bergenstråhle, A. Jurek, J. F. Navarro and J. Lundeberg (2020). “Single-cell and spatial transcriptomics enables probabilistic inference of cell type topography.” Communications biology 3(1): 1–8.

Andersson, A., J. Bergenstråhle, M. Asp, L. Bergenstråhle, A. Jurek, J. F. Navarro and J. J. C. b. Lundeberg (2020). “Single-cell and spatial transcriptomics enables probabilistic inference of cell type topography.” 3(1): 1–8.

Angermueller, C., H. J. Lee, W. Reik and O. Stegle (2017). “DeepCpG: accurate prediction of single-cell DNA methylation states using deep learning.” Genome biology 18(1): 1–13.

Arisdakessian, C., O. Poirion, B. Yunits, X. Zhu and L. X. Garmire (2019). “DeepImpute: an accurate, fast, and scalable deep neural network method to impute single-cell RNA-seq data.” Genome biology 20(1): 1–14.

Arjovsky, M., S. Chintala and L. Bottou (2017). Wasserstein generative adversarial networks. International conference on machine learning, PMLR.

Armacki, A. (2018). Application of Autoencoders on Single-cell Data, University OF Novi Sad.

Atta, L. and J. Fan (2021). “VeloViz: RNA-velocity informed 2D embeddings for visualizing cellular trajectories.” BioRxiv.

Azizi, E., A. J. Carr, G. Plitas, A. E. Cornish, C. Konopacki, S. Prabhakaran, J. Nainys, K. Wu, V. Kiseliovas and M. Setty (2018). “Single-cell map of diverse immune phenotypes in the breast tumor microenvironment.” Cell 174(5): 1293–1308. e1236.

Bacher, R., L.-F. Chu, N. Leng, A. P. Gasch, J. A. Thomson, R. M. Stewart, M. Newton and C. Kendziorski (2017). “SCnorm: robust normalization of single-cell RNA-seq data.” Nature methods 14(6): 584.

Badsha, M. B., R. Li, B. Liu, Y. I. Li, M. Xian, N. E. Banovich and A. Qiuyan (2020). “Imputation of single-cell gene expression with an autoencoder neural network Running title: Autoencoder for imputation of single-cell gene expression.” Quantitative Biology 8(1): 78–94.

Baek, S. and I. Lee (2020). “Single-cell ATAC sequencing analysis: From data preprocessing to hypothesis generation.” Computational and structural biotechnology journal.

Bahdanau, D., J. Chorowski, D. Serdyuk, P. Brakel and Y. Bengio (2016). End-to-end attention-based large vocabulary speech recognition. 2016 IEEE international conference on acoustics, speech and signal processing (ICASSP), IEEE.

Becht, E., L. McInnes, J. Healy, C.-A. Dutertre, I. W. Kwok, L. G. Ng, F. Ginhoux and E. W. Newell (2019). “Dimensionality reduction for visualizing single-cell data using UMAP.” Nature biotechnology 37(1): 38–44.

Bergen, V., M. Lange, S. Peidli, F. A. Wolf and F. J. Theis (2020). “Generalizing RNA velocity to transient cell states through dynamical modeling.” Nature biotechnology 38(12): 1408–1414.

Buettner, F., K. N. Natarajan, F. P. Casale, V. Proserpio, A. Scialdone, F. J. Theis, S. A. Teichmann, J. C. Marioni and O. Stegle (2015). “Computational analysis of cell-to-cell heterogeneity in single-cell RNA-sequencing data reveals hidden subpopulations of cells.” Nature biotechnology 33(2): 155–160.

Buettner, F. and F. J. Theis (2012). “A novel approach for resolving differences in single-cell gene expression patterns from zygote to blastocyst.” Bioinformatics 28(18): i626–i632.

Büttner, M., Z. Miao, F. A. Wolf, S. A. Teichmann and F. J. Theis (2019). “A test metric for assessing single-cell RNA-seq batch correction.” Nature methods 16(1): 43–49.

Cao, J., M. Spielmann, X. Qiu, X. Huang, D. M. Ibrahim, A. J. Hill, F. Zhang, S. Mundlos, L. Christiansen and F. Steemers (2019). “The single-cell transcriptional landscape of mammalian organogenesis.” Nature 566(7745): 496–502.

Cao, K., X. Bai, Y. Hong and L. Wan (2020). “Unsupervised topological alignment for single-cell multi-omics integration.” Bioinformatics 36(Supplement_1): i48–i56.

Cao, Y., L. Fu, J. Wu, Q. Peng, Q. Nie, J. Zhang and X. Xie (2021). “SAILER: Scalable and Accurate Invariant Representation Learning for Single-Cell ATAC-Seq Processing and Integration.” BioRxiv.

Chellapilla, K., S. Puri and P. Simard (2006). High performance convolutional neural networks for document processing. Tenth International Workshop on Frontiers in Handwriting Recognition, Suvisoft.

Chen, L., W. Wang, Y. Zhai and M. Deng (2020). “Deep soft K-means clustering with self-training for single-cell RNA sequence data.” NAR Genomics and Bioinformatics 2(2): lqaa039.

Davie, K., J. Janssens, D. Koldere, M. De Waegeneer, U. Pech, Ł. Kreft, S. Aibar, S. Makhzami, V. Christiaens and C. B. González-Blas (2018). “A single-cell transcriptome atlas of the aging Drosophila brain.” Cell 174(4): 982–998. e920.

Deng, Y., F. Bao, Q. Dai, L. F. Wu and S. J. Altschuler (2019). “Scalable analysis of cell-type composition from single-cell transcriptomics using deep recurrent learning.” Nature methods 16(4): 311–314.

Ding, B., L. Zheng, Y. Zhu, N. Li, H. Jia, R. Ai, A. Wildberg and W. Wang (2015). “Normalization and noise reduction for single cell RNA-seq experiments.” Bioinformatics 31(13): 2225–2227.

Ding, J., A. Condon and S. P. Shah (2018). “Interpretable dimensionality reduction of single cell transcriptome data with deep generative models.” Nature communications 9(1): 1–13.

Dziugaite, G. K., D. M. Roy and Z. Ghahramani (2015). “Training generative neural networks via maximum mean discrepancy optimization.” arXiv preprint 1505.03906.

Elosua-Bayes, M., P. Nieto, E. Mereu, I. Gut and H. Heyn (2021). “SPOTlight: seeded NMF regression to deconvolute spatial transcriptomics spots with single-cell transcriptomes.” Nucleic acids research 49(9): e50–e50.

Engel, J., K. K. Agrawal, S. Chen, I. Gulrajani, C. Donahue and A. Roberts (2019). “Gansynth: Adversarial neural audio synthesis.” arXiv preprint 1902.08710.

Eraslan, G., L. M. Simon, M. Mircea, N. S. Mueller and F. J. Theis (2019). “Single-cell RNA-seq denoising using a deep count autoencoder.” Nature communications 10(1): 1–14.

Esteban, C., S. L. Hyland and G. Rätsch (2017). “Real-valued (medical) time series generation with recurrent conditional gans.” arXiv preprint 1706.02633.

Fedus, W., I. Goodfellow and A. M. Dai (2018). “Maskgan: better text generation via filling in the_.” arXiv preprint 1801.07736.

Fei, T. and T. Yu (2020). “scBatch: batch-effect correction of RNA-seq data through sample distance matrix adjustment.” Bioinformatics.

Finak, G., A. McDavid, M. Yajima, J. Deng, V. Gersuk, A. K. Shalek, C. K. Slichter, H. W. Miller, M. J. McElrath and M. Prlic (2015). “MAST: a flexible statistical framework for assessing transcriptional changes and characterizing heterogeneity in single-cell RNA sequencing data.” Genome biology 16(1): 1–13.

Fritzsch, F. S., C. Dusny, O. Frick and A. Schmid (2012). “Single-cell analysis in biotechnology, systems biology, and biocatalysis.” Annual review of chemical biomolecular engineering 3: 129–155.

Fu, L., L. Zhang, E. Dollinger, Q. Peng, Q. Nie and X. Xie (2020). “Predicting transcription factor binding in single cells through deep learning.” Science Advances 6(51): eaba9031.

Furey, T. S. (2012). “ChIP–seq and beyond: new and improved methodologies to detect and characterize protein–DNA interactions.” Nature Reviews Genetics 13(12): 840–852.

Girshick, R., J. Donahue, T. Darrell and J. Malik (2014). Rich feature hierarchies for accurate object detection and semantic segmentation. Proceedings of the IEEE conference on computer vision and pattern recognition.

Gong, W., I.-Y. Kwak, P. Pota, N. Koyano-Nakagawa and D. J. Garry (2018). “DrImpute: imputing dropout events in single cell RNA sequencing data.” BMC bioinformatics 19(1): 220.

Goodfellow, I., Y. Bengio, A. Courville and Y. Bengio (2016). Deep learning, MIT press Cambridge.

Goodfellow, I. J., J. Pouget-Abadie, M. Mirza, B. Xu, D. Warde-Farley, S. Ozair, A. Courville and Y. Bengio (2014). “Generative adversarial networks.” arXiv preprint 1406.2661.

Grapov, D., J. Fahrmann, K. Wanichthanarak and S. Khoomrung (2018). “Rise of deep learning for genomic, proteomic, and metabolomic data integration in precision medicine.” Omics: a journal of integrative biology 22(10): 630–636.

Griffiths, J. A., A. Scialdone and J. C. Marioni (2018). “Using single-cell genomics to understand developmental processes and cell fate decisions.” Molecular systems biology 14(4): e8046.

Grønbech, C. H., M. F. Vording, P. N. Timshel, C. K. Sønderby, T. H. Pers and O. Winther (2020). “scVAE: Variational auto-encoders for single-cell gene expression data.” Bioinformatics 36(16): 4415–4422.

Guo, X., E. Zhu, X. Liu and J. Yin (2018). Deep embedded clustering with data augmentation. Asian conference on machine learning.

Haber, A. L., M. Biton, N. Rogel, R. H. Herbst, K. Shekhar, C. Smillie, G. Burgin, T. M. Delorey, M. R. Howitt and Y. Katz (2017). “A single-cell survey of the small intestinal epithelium.” Nature 551(7680): 333–339.

Haghverdi, L., F. Buettner and F. J. Theis (2015). “Diffusion maps for high-dimensional single-cell analysis of differentiation data.” Bioinformatics 31(18): 2989–2998.

Haghverdi, L., A. T. Lun, M. D. Morgan and J. C. Marioni (2018). “Batch effects in single-cell RNA-sequencing data are corrected by matching mutual nearest neighbors.” Nature biotechnology 36(5): 421–427.

Han, R. H. and X. T. Zhang (2021). “AImmune: a new blood-based machine learning approach to improving immune profiling analysis on COVID-19 patients.” medRxiv: 2021.2011.2026.21266883.

Han, X., R. Wang, Y. Zhou, L. Fei, H. Sun, S. Lai, A. Saadatpour, Z. Zhou, H. Chen and F. Ye (2018). “Mapping the mouse cell atlas by microwell-seq.” Cell 172(5): 1091–1107. e1017.

Han, X., Z. Zhou, L. Fei, H. Sun, R. Wang, Y. Chen, H. Chen, J. Wang, H. Tang and W. Ge (2020). “Construction of a human cell landscape at single-cell level.” Nature 581(7808): 303–309.

Hannun, A., C. Case, J. Casper, B. Catanzaro, G. Diamos, E. Elsen, R. Prenger, S. Satheesh, S. Sengupta and A. Coates (2014). “Deep speech: Scaling up end-to-end speech recognition.” arXiv preprint 1412.5567.

Hao, Y., S. Hao, E. Andersen-Nissen, W. M. Mauck, S. Zheng, A. Butler, M. J. Lee, A. J. Wilk, C. Darby, M. Zager, P. Hoffman, M. Stoeckius, E. Papalexi, E. P. Mimitou, J. Jain, A. Srivastava, T. Stuart, L. M. Fleming, B. Yeung, A. J. Rogers, J. M. McElrath, C. A. Blish, R. Gottardo, P. Smibert and R. Satija (2021). “Integrated analysis of multimodal single-cell data.” Cell 184(13): 3573–3587.e3529.

He, J., D. Spokoyny, G. Neubig and T. Berg-Kirkpatrick (2019). “Lagging inference networks and posterior collapse in variational autoencoders.” arXiv preprint 1901.05534.

He, K., X. Zhang, S. Ren and J. Sun (2016). Deep residual learning for image recognition. Proceedings of the IEEE conference on computer vision and pattern recognition.

Hentges, L. D., M. D. Sergeant, D. J. Downes, J. R. Hughes and S. Taylor (2021). “LanceOtron: a deep learning peak caller for ATAC-seq, ChIP-seq, and DNase-seq.” BioRxiv.

Heydari, A. A., O. A. Davalos, L. Zhao, K. K. Hoyer and S. S. Sindi (2021). “ACTIVA: realistic single-cell RNA-seq generation with automatic cell-type identification using introspective variational autoencoders.” bioRxiv 2021.01.28.428725; doi: https://doi.org/10.1101/2021.01.28.428725.

Heydari, A. A. and A. Mehmood (2020). SRVAE: super resolution using variational autoencoders. Pattern Recognition and Tracking XXXI, International Society for Optics and Photonics.

Heydari, A. A., C. A. Thompson and A. Mehmood (2019). “Softadapt: Techniques for adaptive loss weighting of neural networks with multi-part loss functions.” 1912.12355; https://arxiv.org/abs/1912.12355.

Hie, B., B. Bryson and B. Berger (2019). “Efficient integration of heterogeneous single-cell transcriptomes using Scanorama.” Nature biotechnology 37(6): 685–691.

Hogan, S. A., A. Courtier, P. F. Cheng, N. F. Jaberg-Bentele, S. M. Goldinger, M. Manuel, S. Perez, N. Plantier, J.-F. Mouret and T. D. L. Nguyen-Kim (2019). “Peripheral blood TCR repertoire profiling may facilitate patient stratification for immunotherapy against melanoma.” Cancer immunology research 7(1): 77–85.

Hosokawa, M., Y. Nishikawa, M. Kogawa and H. Takeyama (2017). “Massively parallel whole genome amplification for single-cell sequencing using droplet microfluidics.” Scientific reports 7(1): 1–11.

Huang, H., Z. Li, R. He, Z. Sun and T. Tan (2018). “Introvae: Introspective variational autoencoders for photographic image synthesis.” arXiv preprint 1807.06358.

Huang, M., J. Wang, E. Torre, H. Dueck, S. Shaffer, R. Bonasio, J. I. Murray, A. Raj, M. Li and N. R. Zhang (2018). “SAVER: gene expression recovery for single-cell RNA sequencing.” Nature methods 15(7): 539–542.

Hubel, D. H. and T. N. Wiesel (1962). “Receptive fields, binocular interaction and functional architecture in the cat’s visual cortex.” The Journal of physiology 160(1): 106–154.

Jeong, H.-H., J. Jia, Y. Dai, L. M. Simon and Z. Zhao (2021). “Investigating Cellular Trajectories in the Severity of COVID-19 and Their Transcriptional Programs Using Machine Learning Approaches.” Genes 12(5): 635.

Jiang, P., J. A. Thomson and R. Stewart (2016). “Quality control of single-cell RNA-seq by SinQC.” Bioinformatics 32(16): 2514–2516.

Jin, W., J. M. Stokes, R. T. Eastman, Z. Itkin, A. V. Zakharov, J. J. Collins, T. S. Jaakkola and R. Barzilay (2021). “Deep learning identifies synergistic drug combinations for treating COVID-19.” Proceedings of the National Academy of Sciences 118(39).

Johnson, W. E., C. Li and A. Rabinovic (2007). “Adjusting batch effects in microarray expression data using empirical Bayes methods.” Biostatistics 8(1): 118–127.

Kharchenko, P. V., L. Silberstein and D. T. J. N. m. Scadden (2014). “Bayesian approach to single-cell differential expression analysis.” 11(7): 740–742.

Kimmel, J. C., A. S. Brack and W. F. Marshall (2019). “Deep convolutional and recurrent neural networks for cell motility discrimination and prediction.” BioRxiv: 159202.

Kingma, D. P. and J. Ba (2015). “Adam: A Method for Stochastic Optimization.” arXiv preprint.

Kingma, D. P. and M. Welling (2013). “Auto-encoding variational bayes.” arXiv preprint 1312.6114.

Kingma, D. P. and M. Welling (2019). “An introduction to variational autoencoders.” arXiv preprint 1906.02691.

Klein, A. M., L. Mazutis, I. Akartuna, N. Tallapragada, A. Veres, V. Li, L. Peshkin, D. A. Weitz and M. W. Kirschner (2015). “Droplet barcoding for single-cell transcriptomics applied to embryonic stem cells.” Cell 161(5): 1187–1201.

Kleshchevnikov, V., A. Shmatko, E. Dann, A. Aivazidis, H. W. King, T. Li, A. Lomakin, V. Kedlian, M. S. Jain, J. S. Park, L. Ramona, E. Tuck, A. Arutyunyan, R. Vento-Tormo, M. Gerstung, L. James, O. Stegle and O. A. Bayraktar (2020). “Comprehensive mapping of tissue cell architecture via integrated single cell and spatial transcriptomics.” 2020.2011.2015.378125.

Krizhevsky, A., I. Sutskever, G. E. Hinton, F. Pereira, C. Burges, L. Bottou and K. Weinberger (2012). “Advances in neural information processing systems.”

Lafzi, A., C. Moutinho, S. Picelli and H. Heyn (2018). “Tutorial: guidelines for the experimental design of single-cell RNA sequencing studies.” Nature protocols 13(12): 2742–2757.

Lähnemann, D., J. Köster, E. Szczurek, D. J. McCarthy, S. C. Hicks, M. D. Robinson, C. A. Vallejos, K. R. Campbell, N. Beerenwinkel and A. Mahfouz (2020). “Eleven grand challenges in single-cell data science.” Genome biology 21(1): 1–35.

Larsen, A. B. L., S. K. Sønderby, H. Larochelle and O. Winther (2016). Autoencoding beyond pixels using a learned similarity metric. International conference on machine learning, PMLR.

LeCun, Y. and Y. Bengio (1995). “Convolutional networks for images, speech, and time series.” The handbook of brain theory and neural networks 3361(10): 1995.

LeCun, Y., Y. Bengio and G. Hinton (2015). “Deep learning.” nature 521(7553): 436–444.

Li, H., U. Shaham, K. P. Stanton, Y. Yao, R. R. Montgomery and Y. Kluger (2017). “Gating mass cytometry data by deep learning.” Bioinformatics 33(21): 3423–3430.

Li, W. V. and J. J. Li (2018). “An accurate and robust imputation method scImpute for single-cell RNA-seq data.” Nature communications 9(1): 1–9.

Li, X., K. Wang, Y. Lyu, H. Pan, J. Zhang, D. Stambolian, K. Susztak, M. P. Reilly, G. Hu and M. Li (2020). “Deep learning enables accurate clustering with batch effect removal in single-cell RNA-seq analysis.” Nature communications 11(1): 1–14.

Lin, K. L., T. Yang, H. Y. Zou, Y. F. Li and C. Z. Huang (2019). “Graphitic C3N4 nanosheet and hemin/G-quadruplex DNAzyme-based label-free chemiluminescence aptasensing for biomarkers.” Talanta 192: 400–406.

Lin, P., M. Troup and J. W. Ho (2017). “CIDR: Ultrafast and accurate clustering through imputation for single-cell RNA-seq data.” Genome biology 18(1): 59.

Lin, S., Y. Lin, J. R. Nery, M. A. Urich, A. Breschi, C. A. Davis, A. Dobin, C. Zaleski, M. A. Beer and W. C. Chapman (2014). “Comparison of the transcriptional landscapes between human and mouse tissues.” Proceedings of the National Academy of Sciences 111(48): 17224–17229.

Lin, Y., T.-Y. Wu, S. Wan, J. Y. Yang, Y. R. Wang and W. H. Wong (2021). “scJoint: transfer learning for data integration of single -cell RNA-seq and ATAC-seq.” BioRxiv: 2020.2012. 2031.424916.

Linker, S. M., L. Urban, S. J. Clark, M. Chhatriwala, S. Amatya, D. J. McCarthy, I. Ebersberger, L. Vallier, W. Reik and O. Stegle (2019). “Combined single-cell profiling of expression and DNA methylation reveals splicing regulation and heterogeneity.” Genome biology 20(1): 1–14.

Liu, J., Y. Huang, R. Singh, J.-P. Vert and W. S. Noble (2019). “Jointly embedding multiple single-cell omics measurements.” BioRxiv: 644310.

Long, J., E. Shelhamer and T. Darrell (2015). Fully convolutional networks for semantic segmentation. Proceedings of the IEEE conference on computer vision and pattern recognition.

Lopez, R., J. Regier, M. B. Cole, M. I. Jordan and N. Yosef (2018). “Deep generative modeling for single-cell transcriptomics.” Nature methods 15(12): 1053–1058.

Love, M. I., W. Huber and S. Anders (2014). “Moderated estimation of fold change and dispersion for RNA-seq data with DESeq2.” Genome biology 15(12): 1–21.

Lucas, J., G. Tucker, R. B. Grosse and M. Norouzi (2019). “Don’t blame the Elbo! a linear Vae perspective on posterior collapse.” Advances in Neural Information Processing Systems 32: 9408–9418.

Luecken, M. D. and F. J. Theis (2019). “Current best practices in single-cell RNA-seq analysis: a tutorial.” Molecular systems biology 15(6): e8746.

Lun, A. T., K. Bach and J. C. Marioni (2016). “Pooling across cells to normalize single-cell RNA sequencing data with many zero counts.” Genome biology 17(1): 1–14.

Luo, X. and Y. Wei (2019). “Batch effects correction with unknown subtypes.” Journal of the American Statistical Association 114(526): 581–594.

Marouf, M., P. Machart, V. Bansal, C. Kilian, D. S. Magruder, C. F. Krebs and S. Bonn (2020). “Realistic in silico generation and augmentation of single-cell RNA-seq data using generative adversarial networks.” Nature communications 11(1): 1–12.

Märtens, K. and C. Yau (2020). “BasisVAE: Translation-invariant feature-level clustering with Variational Autoencoders.” arXiv preprint 2003.03462.

Marx, V. (2021). “Method of the Year: spatially resolved transcriptomics.” Nature Methods 18(1): 9–14.

Metz, L., B. Poole, D. Pfau and J. Sohl-Dickstein (2016). “Unrolled generative adversarial networks.” arXiv preprint 1611.02163.

Mongia, A., D. Sengupta and A. Majumdar (2020). “deepmc: Deep matrix completion for imputation of single-cell rna-seq data.” Journal of Computational Biology 27(7): 1011–1019.

Montoro, D. T., A. L. Haber, M. Biton, V. Vinarsky, B. Lin, S. E. Birket, F. Yuan, S. Chen, H. M. Leung and J. Villoria (2018). “A revised airway epithelial hierarchy includes CFTR-expressing ionocytes.” Nature 560(7718): 319–324.

Moussa, M. and I. I. Măndoiu (2019). “Locality sensitive imputation for single cell RNA-seq data.” Journal of Computational Biology 26(8): 822–835.

Nair, V. and G. E. Hinton (2010). Rectified linear units improve restricted boltzmann machines. Icml.

Parekh, S., C. Ziegenhain, B. Vieth, W. Enard and I. Hellmann (2018). “zUMIs-a fast and flexible pipeline to process RNA sequencing data with UMIs.” Gigascience.

Patruno, L., D. Maspero, F. Craighero, F. Angaroni, M. Antoniotti and A. Graudenzi (2020). “A review of computational strategies for denoising and imputation of single-cell transcriptomic data.” Briefings in Bioinformatics 22(4).

Pearson, K. (1901). “LIII. On lines and planes of closest fit to systems of points in space.” The London, Edinburgh, Dublin philosophical magazine journal of science 2(11): 559–572.

Peng, J., X. Wang and X. Shang (2019). “Combining gene ontology with deep neural networks to enhance the clustering of single cell RNA-Seq data.” BMC bioinformatics 20(8): 284.

Peng, T., G. Chen and K. Tan (2021). “GLUER: integrative analysis of single-cell omics and imaging data by deep neural network.” BioRxiv.

Pham, D. T., X. Tan, J. Xu, L. F. Grice, P. Y. Lam, A. Raghubar, J. Vukovic, M. J. Ruitenberg and Q. H. Nguyen (2020). “stLearn: integrating spatial location, tissue morphology and gene expression to find cell types, cell-cell interactions and spatial trajectories within undissociated tissues.” BioRxiv.

Pierson, E. and C. J. G. b. Yau (2015). “ZIFA: Dimensionality reduction for zero-inflated single-cell gene expression analysis.” 16(1): 1–10.

Plass, M., J. Solana, F. A. Wolf, S. Ayoub, A. Misios, P. Glažar, B. Obermayer, F. J. Theis, C. Kocks and N. Rajewsky (2018). “Cell type atlas and lineage tree of a whole complex animal by single-cell transcriptomics.” Science 360(6391).

Polański, K., M. D. Young, Z. Miao, K. B. Meyer, S. A. Teichmann and J.-E. Park (2020). “BBKNN: fast batch alignment of single cell transcriptomes.” Bioinformatics 36(3): 964–965.

Qiu, P. (2020). “Embracing the dropouts in single-cell RNA-seq analysis.” Nature communications 11(1): 1–9.

Ran, D., S. Zhang, N. Lytal and L. An (2019). “scDoc: Correcting Drop-out Events in Single-cell RNA-seq Data.” BioRxiv: 731638.

Regev, A., S. A. Teichmann, E. S. Lander, I. Amit, C. Benoist, E. Birney, B. Bodenmiller, P. Campbell, P. Carninci and M. Clatworthy (2017). “Science forum: the human cell atlas.” Elife 6: e27041.

Ritchie, M. E., B. Phipson, D. Wu, Y. Hu, C. W. Law, W. Shi and G. K. Smyth (2015). “limma powers differential expression analyses for RNA-sequencing and microarray studies.” Nucleic acids research 43(7): e47–e47.

Rodriques, S. G., R. R. Stickels, A. Goeva, C. A. Martin, E. Murray, C. R. Vanderburg, J. Welch, L. M. Chen, F. Chen and E. Z. Macosko (2019). “Slide-seq: A scalable technology for measuring genome-wide expression at high spatial resolution.” Science 363(6434): 1463–1467.

Roth, A., J. Khattra, D. Yap, A. Wan, E. Laks, J. Biele, G. Ha, S. Aparicio, A. Bouchard-Côté and S. P. Shah (2014). “PyClone: statistical inference of clonal population structure in cancer.” Nature methods 11(4): 396–398.

Rozenblatt-Rosen, O., M. J. Stubbington, A. Regev and S. A. Teichmann (2017). “The Human Cell Atlas: from vision to reality.” Nature News 550(7677): 451.

Rumelhart, D. E., G. E. Hinton and R. J. Williams (1986). “Learning representations by back-propagating errors.” nature 323(6088): 533–536.

Saha, I., N. Ghosh, D. Maity, A. Seal and D. Plewczynski (2021). “COVID-DeepPredictor: Recurrent Neural Network to Predict SARS-CoV-2 and Other Pathogenic Viruses.” Frontiers in genetics 12: 83.

Semeniuta, S., A. Severyn and E. Barth (2017). “A hybrid convolutional variational autoencoder for text generation.” arXiv preprint 1702.02390.

Sengupta, D., N. A. Rayan, M. Lim, B. Lim and S. Prabhakar (2016). “Fast, scalable and accurate differential expression analysis for single cells.” BioRxiv: 049734.

Shaham, U. (2018). “Batch Effect Removal via Batch-Free Encoding.” BioRxiv: 380816.

Shaham, U., K. P. Stanton, J. Zhao, H. Li, K. Raddassi, R. Montgomery and Y. Kluger (2017). “Removal of batch effects using distribution-matching residual networks.” Bioinformatics 33(16): 2539–2546.

Shi, S., Q. Wang, P. Xu and X. Chu (2016). Benchmarking state-of-the-art deep learning software tools. 2016 7th International Conference on Cloud Computing and Big Data (CCBD), IEEE.

Shorten, C. and T. M. Khoshgoftaar (2019). “A survey on image data augmentation for deep learning.” Journal of Big Data 6(1): 1–48.

Ståhl, P. L., F. Salmén, S. Vickovic, A. Lundmark, J. F. Navarro, J. Magnusson, S. Giacomello, M. Asp, J. O. Westholm and M. Huss (2016). “Visualization and analysis of gene expression in tissue sections by spatial transcriptomics.” Science 353(6294): 78–82.

Stark, S. G., J. Ficek, F. Locatello, X. Bonilla, S. Chevrier, F. Singer, G. Rätsch and K.-V. Lehmann (2020). “SCIM: universal single-cell matching with unpaired feature sets.” Bioinformatics 36(Supplement_2): i919–i927.

Steinkraus, D., I. Buck and P. Simard (2005). Using GPUs for machine learning algorithms. Eighth International Conference on Document Analysis and Recognition (ICDAR’05), IEEE.

Stuart, T., A. Butler, P. Hoffman, C. Hafemeister, E. Papalexi, W. M. Mauck III, Y. Hao, M. Stoeckius, P. Smibert and R. Satija (2019). “Comprehensive integration of single-cell data.” Cell 177(7): 1888–1902. e1821.

Sun, Y., D. Liang, X. Wang and X. Tang (2015). “Deepid3: Face recognition with very deep neural networks.” arXiv preprint 1502.00873.

Tabula Muris, C., c. Overall, c. Logistical, c. Organ, processing, p. Library, sequencing, a. Computational data, a. Cell type, g. Writing, g. Supplemental text writing and i. Principal (2018). “Single-cell transcriptomics of 20 mouse organs creates a Tabula Muris.” Nature 562(7727): 367–372.

Talwar, D., A. Mongia, D. Sengupta and A. Majumdar (2018). “AutoImpute: Autoencoder based imputation of single-cell RNA-seq data.” Scientific reports 8(1): 1–11.

Teichmann, S. and M. Efremova (2020). “Method of the Year 2019: single-cell multimodal omics.” Nat. Methods 17(1).

Thibodeau, A., S. Khetan, A. Eroglu, R. Tewhey, M. L. Stitzel and D. Ucar (2020). “CoRE-ATAC: A Deep Learning model for the functional Classification of Regulatory Elements from single cell and bulk ATAC-seq data.” BioRxiv.

Tian, T., J. Wan, Q. Song and Z. Wei (2019). “Clustering single-cell RNA-seq data with a model-based deep learning approach.” Nature Machine Intelligence 1(4): 191–198.

Titsias, M. and N. D. Lawrence (2010). Bayesian Gaussian process latent variable model. Proceedings of the Thirteenth International Conference on Artificial Intelligence and Statistics, JMLR Workshop and Conference Proceedings.

Tolstikhin, I., O. Bousquet, S. Gelly and B. Schoelkopf (2017). “Wasserstein auto-encoders.” arXiv preprint 1711.01558.

Tran, H. T. N., K. S. Ang, M. Chevrier, X. Zhang, N. Y. S. Lee, M. Goh and J. Chen (2020). “A benchmark of batch-effect correction methods for single-cell RNA sequencing data.” Genome biology 21(1): 1–32.

Vallejos, C. A., J. C. Marioni and S. Richardson (2015). “BASiCS: Bayesian analysis of single-cell sequencing data.” PLoS computational biology 11(6).

Van der Maaten, L. and G. Hinton (2008). “Visualizing data using t-SNE.” Journal of machine learning research 9(11).

van Dijk, D., J. Nainys, R. Sharma, P. Kaithail, A. J. Carr, K. R. Moon, L. Mazutis, G. Wolf, S. Krishnaswamy and D. Pe’er (2017). “MAGIC: A diffusion-based imputation method reveals gene-gene interactions in single-cell RNA-sequencing data.” BioRxiv: 111591.

Vickovic, S., G. Eraslan, F. Salmén, J. Klughammer, L. Stenbeck, D. Schapiro, T. Äijö, R. Bonneau, L. Bergenstråhle and J. F. Navarro (2019). “High-definition spatial transcriptomics for in situ tissue profiling.” Nature methods 16(10): 987–990.

Vondrick, C., H. Pirsiavash and A. Torralba (2016). “Generating videos with scene dynamics.” arXiv preprint 1609.02612.

Wang, B., D. Ramazzotti, L. De Sano, J. Zhu, E. Pierson and S. Batzoglou (2017). “SIMLR: a tool for large-scale single-cell analysis by multi-kernel learning.” BioRxiv: 118901.

Wang, D. and J. Gu (2018). “VASC: dimension reduction and visualization of single-cell RNA-seq data by deep variational autoencoder.” Genomics, proteomics & bioinformatics 16(5): 320–331.

Wang, T., T. S. Johnson, W. Shao, Z. Lu, B. R. Helm, J. Zhang and K. Huang (2019). “BERMUDA: a novel deep transfer learning method for single-cell RNA sequencing batch correction reveals hidden high-resolution cellular subtypes.” Genome biology 20(1): 1–15.

Wang, Z., Q. She and T. E. Ward (2021). “Generative adversarial networks in computer vision: A survey and taxonomy.” ACM Computing Surveys (CSUR) 54(2): 1–38.

Wani, N. and K. Raza (2019). “Integrative approaches to reconstruct regulatory networks from multi-omics data: a review of state-of-the-art methods.” Computational biology and chemistry 83: 107120.

Wu, Y., M. Schuster, Z. Chen, Q. V. Le, M. Norouzi, W. Macherey, M. Krikun, Y. Cao, Q. Gao and K. Macherey (2016). “Google’s neural machine translation system: Bridging the gap between human and machine translation.” arXiv preprint 1609.08144.

Xiong, L., K. Xu, K. Tian, Y. Shao, L. Tang, G. Gao, M. Zhang, T. Jiang and Q. C. Zhang (2019). “SCALE method for single -cell ATAC-seq analysis via latent feature extraction.” Nature communications 10(1): 1–10.

Xu, Y., P. Das and R. P. McCord (2021). “SMILE: Mutual Information Learning for Integration of Single Cell Omics Data.” BioRxiv.

Xu, Y., Z. Zhang, L. You, J. Liu, Z. Fan and X. Zhou (2020). “scIGANs: single-cell RNA-seq imputation using generative adversarial networks.” Nucleic Acids Research 48(15): e85–e85.

Yan, F., D. R. Powell, D. J. Curtis and N. C. Wong (2020). “From reads to insight: a hitchhiker’s guide to ATAC-seq data analysis.” Genome biology 21(1): 22.

Yang, L., J. Liu, Q. Lu, A. D. Riggs and X. Wu (2017). “SAIC: an iterative clustering approach for analysis of single cell RNA-seq data.” BMC genomics 18(6): 689.

Yang, X., Y.-N. Chen, D. Hakkani-Tür, P. Crook, X. Li, J. Gao and L. Deng (2017). End-to-end joint learning of natural language understanding and dialogue manager. 2017 IEEE International Conference on Acoustics, Speech and Signal Processing (ICASSP), IEEE.

Yang, Z., Z. Hu, R. Salakhutdinov and T. Berg-Kirkpatrick (2017). Improved variational autoencoders for text modeling using dilated convolutions. International conference on machine learning, PMLR.

Yuan, Y. and Z. Bar-Joseph (2019). “Deep learning for inferring gene relationships from single-cell expression data.” Proceedings of the National Academy of Sciences 116(52): 27151–27158.

Zafar, H., N. Navin, K. Chen and L. Nakhleh (2019). “SiCloneFit: Bayesian inference of population structure, genotype, and phylogeny of tumor clones from single-cell genome sequencing data.” Genome research 29(11): 1847–1859.

Zappia, L., B. Phipson and A. Oshlack (2017). “Splatter: simulation of single-cell RNA sequencing data.” Genome biology 18(1): 174.

Zhang, F., Y. Wu and W. Tian (2019). “A novel approach to remove the batch effect of single-cell data.” Cell discovery 5(1): 1–4.

Zhang, Z., F. Cui, C. Lin, L. Zhao, C. Wang and Q. Zou (2021). “Critical downstream analysis steps for single-cell RNA sequencing data.” Briefings in Bioinformatics.

Zhao, S., J. Song and S. Ermon (2017). “Infovae: Information maximizing variational autoencoders.” arXiv preprint 1706.02262.

Zhao, Z.-Q., P. Zheng, S.-t. Xu and X. Wu (2019). “Object detection with deep learning: A review.” IEEE transactions on neural networks and learning systems 30(11): 3212–3232.

Zheng, G. X., J. M. Terry, P. Belgrader, P. Ryvkin, Z. W. Bent, R. Wilson, S. B. Ziraldo, T. D. Wheeler, G. P. McDermott and J. Zhu (2017). “Massively parallel digital transcriptional profiling of single cells.” Nature communications 8(1): 1–12.

Zheng, J. and K. Wang (2019). “Emerging deep learning methods for single-cell RNA-seq data analysis.” Quantitative Biology 7(4): 247–254.

Zheng, R., M. Li, Z. Liang, F.-X. Wu, Y. Pan and J. Wang (2019). “SinNLRR: a robust subspace clustering method for cell type detection by non-negative and low-rank representation.” Bioinformatics 35(19): 3642–3650.

Zhou, J. and O. G. Troyanskaya (2015). “Predicting effects of noncoding variants with deep learning–based sequence model.” Nature methods 12(10): 931–934.

Zhu, C., S. Preissl and B. Ren (2020). “Single-cell multimodal omics: the power of many.” Nature methods 17(1): 11–14.

Zhu, J.-Y., P. Krähenbühl, E. Shechtman and A. A. Efros (2016). Generative visual manipulation on the natural image manifold. European conference on computer vision, Springer.

Zilionis, R., J. Nainys, A. Veres, V. Savova, D. Zemmour, A. M. Klein and L. Mazutis (2017). “Single-cell barcoding and sequencing using droplet microfluidics.” Nature protocols 12(1): 44–73.

Zuo, C. and L. Chen (2020). “Deep-joint-learning analysis model of single cell transcriptome and open chromatin accessibility data.” Briefings in Bioinformatics 22(4).

